# Ran-GTP is non-essential to activate NuMA for spindle pole focusing, but dynamically polarizes HURP to control mitotic spindle length

**DOI:** 10.1101/473538

**Authors:** Kenta Tsuchiya, Hisato Hayashi, Momoko Nishina, Masako Okumura, Yoshikatsu Sato, Masato T. Kanemaki, Gohta Goshima, Tomomi Kiyomitsu

**Affiliations:** Division of Biological Science, Graduate School of Science, Nagoya University, Chikusa-ku, Nagoya 464-8602, Japan; Precursory Research for Embryonic Science and Technology (PRESTO) Program, Japan Science and Technology Agency, 4-1-8 Honcho Kawaguchi, Saitama 332-0012, Japan; Department of Chromosome Science, National Institute of Genetics, Research Organization of Information and Systems (ROIS), and Department of Genetics, SOKENDAI (The Graduate University of Advanced Studies), Yata 1111, Mishima, Shizuoka 411-8540, Japan; Okinawa Institute of Science and Technology Graduate University, 1919-1 Tancha, Onna-son, Kunigami-gun, Okinawa 904-0495, Japan

**Author notes:** These authors contributed equally to this work (Kenta Tsuchiya, Hisato Hayashi, Momoko Nishina, Masako Okumura).

## Abstract

During mitosis, a bipolar spindle is assembled around chromosomes to efficiently capture chromosomes. Previous work proposed that a chromosome-derived Ran-GTP gradient promotes spindle assembly around chromosomes by liberating spindle assembly factors (SAFs) from inhibitory importins. However, Ran’s dual functions in interphase nucleocytoplasmic transport and mitotic spindle assembly have made it difficult to assess its mitotic roles in somatic cells. Here, using auxin-inducible degron technology in human cells, we developed acute mitotic degradation assays to dissect Ran’s mitotic roles systematically and separately from its interphase function. In contrast to the prevailing model, we found that the Ran pathway is not essential for spindle assembly activities that occur at sites spatially separated from chromosomes, including activating NuMA for spindle pole focusing or for targeting TPX2. In contrast, Ran-GTP is required to localize HURP and HSET specifically at chromosome-proximal regions. We demonstrated that Ran-GTP and importin-β coordinately promote HURP’s dynamic microtubule binding-dissociation cycle near chromosomes, which results in stable kinetochore-fiber formation. Intriguingly, this pathway acts to establish proper spindle length preferentially during prometaphase, rather than metaphase. Together, we propose that the Ran pathway is required to activate SAFs specifically near chromosomes, but not generally during human mitotic spindle assembly. Ran-dependent spindle assembly is likely coupled with parallel pathways to activate SAFs, including NuMA, for spindle pole focusing away from chromosomes.

**Highlights:**

- Using auxin-inducible degron technology, we developed mitotic degradation assays for the Ran pathway in human cells.
- The Ran pathway is non-essential to activate NuMA for spindle pole focusing.
- The Ran pathway dynamically polarizes HURP and defines mitotic spindle length preferentially during prometaphase.
- Ran-GTP is required to activate SAFs specifically near chromosomes, but not generally, in human mitotic cells.

## Introduction

During cell division, a microtubule-based spindle structure is assembled around chromosomes to efficiently capture and segregate duplicated chromosomes into daughter cells [1, 2]. To assemble a spindle around chromosomes, chromosomes generate a gradient of Ran-GTP, a GTP-bound form of Ran, in animal cells [3, 4]. Ran-GTP is produced by regulator of chromosome condensation 1 (RCC1), a guanine nucleotide exchange factor for Ran [5], and is hydrolyzed to Ran-GDP by RanGAP1, a GTPase-activating protein for Ran [6]. Because RCC1 and RanGAP1 mainly localize on chromosomes and in cytoplasm, respectively, these opposing enzymes create a chromosome-derived Ran-GTP gradient after the nuclear envelope breaks down (Fig. 2A). During interphase, these enzymes generate different Ran-GTP concentrations in the nucleus and cytoplasm, which drives nucleocytoplasmic transport [4]. The Ran-GTP gradient has been best characterized in *Xenopus* egg extracts [7, 8], but is also found in other meiotic and mitotic cell types [9–11]. Recent studies indicate that Ran-GTP is essential for acentrosomal spindle assembly in female meiosis [9, 12, 13], but the significance of Ran-GTP in mitotic spindle assembly has been debated [10, 11, 14]. The dual functions of Ran in both interphase and mitosis have made it difficult to identify its mitotic roles in somatic cells.

As in mechanisms in nucleocytoplasmic transport, Ran-GTP binds to importin-β and releases inhibitory importins from SAFs, thereby activating SAFs near chromosomes (Fig. 2A) [15–18]. Once activated, most SAFs interact with microtubules and spatially regulate microtubule nucleation, dynamics, transport, and cross-linking, to create specialized local structures of the spindle [3, 4]. For instance, nuclear mitotic apparatus protein (NuMA) recognizes minus-ends of microtubules and transports and crosslinks microtubules in cooperation with cytoplasmic dynein, a minus-end-directed motor, to focus spindle microtubules at the poles of mammalian cells [19–22]. The targeting protein for Xklp2 (TPX2) is required for spindle pole organization [23, 24] and stimulates microtubule nucleation in a Ran- and importin-α-regulated manner [25–27]. Kinesin-14 HSET/XCTK2 cross-links both parallel and anti-parallel microtubules near chromosomes, but preferentially cross-links parallel microtubules near the spindle poles [28–30]. Hepatoma upregulated protein (HURP) accumulates on microtubules near chromosomes to form stabilized kinetochore-fibers (k-fibers) [31].

Most SAFs, including NuMA, TPX2, and HSET contain a nuclear localization sequence/signal (NLS) [28, 32, 33]. The NLS is specifically recognized by importin-α, which forms hetero-dimer with importin-β through an importin-β binding (IBB) domain (Fig. 2A). On the other hand, some SAFs, such as HURP, are directly recognized by importin-β (Fig. 2A) [31]. Because SAFs represent a small fraction of NLS-bearing nuclear proteins and need to be regulated effectively and selectively in mitosis, each SAF interacts with importins in specific ways to reduce competition with other nuclear proteins and SAFs [32, 33].

In mitotic human cells, NuMA localizes to the spindle poles and the polar cell cortex, where it facilitates spindle-pole focusing and astral microtubule capture/pulling, respectively [19, 20, 34]. Recent structural and *in vitro* studies have demonstrated that NuMA’s microtubule-binding activities are inhibited by steric blockage of importin-β, mediated by importin-α [32], but this model has not been rigorously tested in cells. In addition, given that the Ran-GTP gradient diminishes with increasing distances from chromosomes, it is unclear whether and how the Ran-GTP gradient activates NuMA at the spindle poles.

To precisely understand mechanisms and significance of Ran-based regulation of SAFs, it is critical to separate Ran’s mitotic roles from its interphase nucleocytoplasmic transport function. To achieve this, we developed mitotic depletion assays for the Ran pathway in human cells by combining mitotic drugs with auxin-inducible degron (AID) technology [35], which allows us to degrade mAID-tag fusion proteins with a half-life of 20 min. In contrast to the prevailing model, we found that degradation of RCC1, RanGAP1, or importin-β does not substantially affect localization and function of NuMA at the spindle poles, even if these proteins were degraded during mitosis. In sharp contrast, the Ran pathway polarizes both HURP and importin-β on k-fibers near chromosomes, where HURP stabilizes k-fibers independently of importin-β. Based on our results, we propose that the Ran-Importin pathway is required to activate SAFs specifically near chromosomes, but not generally, in human mitotic cells.

## Results

### In human cells, NuMA focuses spindle microtubules at spindle poles using its C-terminal conserved microtubule-binding domain

NuMA functions in spindle microtubule focusing in cultured mammalian cells [19–22]. Silk et al. demonstrated that NuMA’s C-terminal microtubule-binding domain (MTBD1) adjacent to a NLS is required for spindle pole focusing in mouse fibroblasts [22] (Fig. 1A). However, this domain is dispensable for spindle pole focusing in mouse keratinocytes [36]. In addition, NuMA has a second microtubule-binding domain (MTBD2) at the C-terminal end (Fig. 1A) [32, 37], which has stronger microtubule-binding activity and is sterically inhibited by importin-β *in vitro* [32]. To understand which domain of NuMA is required for spindle pole focusing in mitotic human cells, we replaced endogenous NuMA with C-terminal truncation mutants in HCT116 cells (Fig. 1A). Endogenous NuMA was visualized by integrating an mAID-mClover-FLAG (mACF) tag into both alleles of the NuMA gene [20]. NuMA-mACF was depleted using the auxin inducible degradation (AID) system following Dox and IAA treatment (S1A) [20, 35], and mCherry-tagged NuMA mutants were simultaneously expressed from the Rosa 26 locus by Dox treatment (Fig. 1B-C, S1B) [20]. Like endogenous NuMA, mCherry-tagged NuMA wild type (WT) accumulated in interphase nuclei (Fig. 1B) and at mitotic spindle poles (Fig. 1C #1) and was able to rescue pole-focusing defects caused by NuMA depletion (Fig. 1C-D #1). NuMA-ΔNLS mutants were unable to localize at nuclei in interphase (Fig. 1B, S1C), but were able to accumulate at spindle poles to rescue pole-focusing defects (Fig. 1C-D #2). As expected, NuMA ΔC-ter mutants, which lack both MTBDs, diffused into the cytoplasm during metaphase (Fig. 1C #5), and were unable to rescue the spindle-pole focusing defect (Fig. 1C-D #5). In contrast, NuMA Δ(NLS+MTBD2) mutants localized around spindle-poles to rescue the focusing defects (Fig. 1C-D #4). However, NuMA Δex24 mutants, which lack NLS and the part of MTBD1 containing the well-conserved NLM motif (Fig. S1D) [38, 39], were unable to fully rescue focusing defects, while localizing around the spindle-poles (Fig. 1C-D #3). These results indicate that NuMA’s MTBD1, but not MTBD2, is essential for spindle pole focusing in human cells.

**Figure 1.**
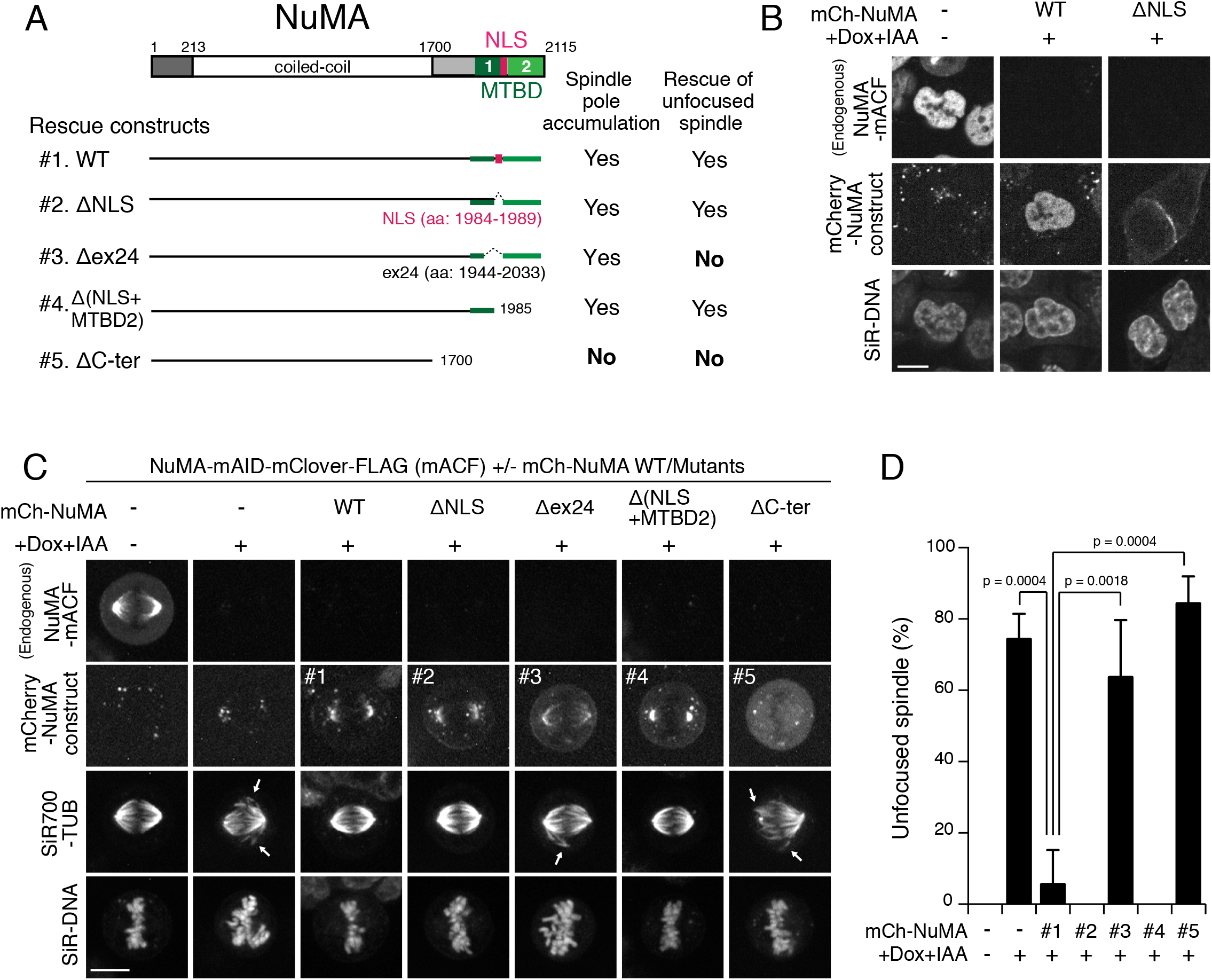
NuMA acts in spindle pole focusing using its conserved microtubule-binding domain in human cells. (A) Full length NuMA and tested NuMA truncation fragments. NLS and a microtubule-binding domain (MTBD) are shown in magenta and green, respectively. (B and C) Interphase (B) and metaphase (C) Metaphase NuMA-mACF cell lines showing live fluorescent images of NuMA-mACF, NuMA-mCh WT or mutants, SiR-DNA and SiR-700 tubulin (TUB) after 24 hr following treatment with Dox and IAA. Arrows in C indicate unfocused microtubules. (D) Quantification of cells with unfocused spindles in each condition from data in (C). Bars indicate means ± SEMs. N = 47 (-/-), 75 (-/+), 31 (#1/+), 30 (#2/+), 31 (#3/+), 30 (#4/+), and 48 (#5/+) from 3 independent experiments. p-values calculated using Dunnett’s multiple comparisons test after one-way ANOVA (*F* (3,6) = 33.81, *p* = 0.0004).

### NuMA localizes at the spindle poles and participates in spindle pole focusing independently of RCC1

NuMA’s MTBD1 is located next to NLS, which is recognized by importin-α [32]. A recent study indicated that the importin-α/β complex sterically inhibits NuMA’s microtubule-binding activity, but is released from NuMA by Ran-GTP *in vitro* [32] (Fig. 2A). To test this model in cells, we next depleted RCC1 (RanGEF) by integrating mAID-mClover (mAC) tag (Fig. 2B, S2A) [35]. In contrast to the model, NuMA accumulated normally around the spindle poles, and spindle microtubules were properly focused in RCC1-depleted cells (Fig. 2B, D, Fig. S2B-C), although metaphase spindle length diminished (Fig. 2B-C), and mitotic duration was slightly delayed (Fig. S2D-F).

To further analyze functions of NuMA in RCC1-depleted cells, we next co-depleted RCC1 and NuMA. Following treatment with Dox and IAA, both RCC1-mAC and NuMA-mAID-mCherry were degraded, and spindle microtubules were not properly focused (Fig. 2D-E, S2G-H). These results indicate that NuMA acts at spindle poles, even in the absence of Ran-GTP in human mitotic cells.

**Figure 2.**
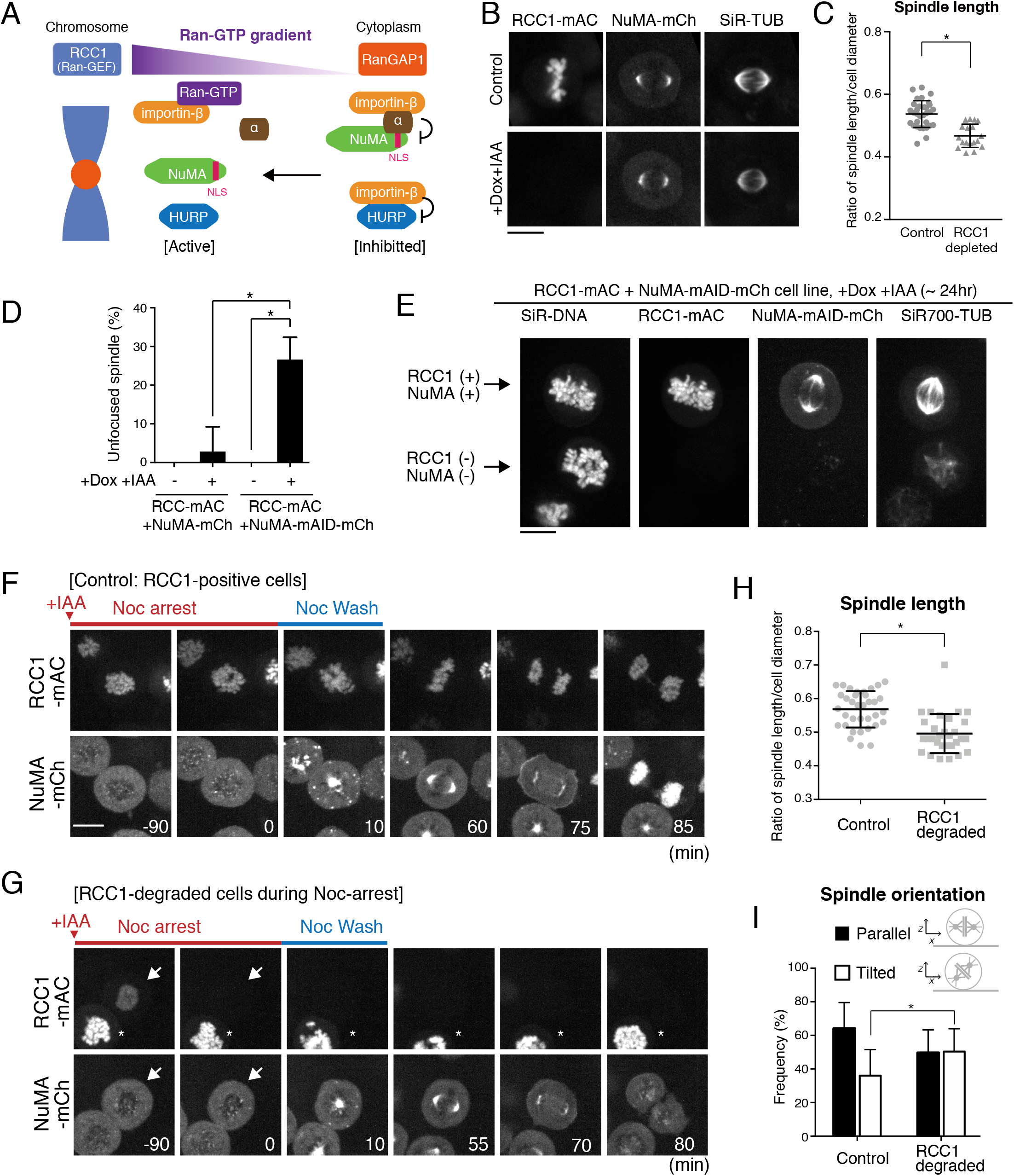
NuMA functions in spindle pole focusing independently of RCC1. (A) The prevailing model of SAF inhibition and activation by importins and Ran-GTP. (B) Metaphase RCC1-mAC cells showing live fluorescent images of RCC1mAC, NuMA-mCherry (mCh), and SiR-TUB after 24 hr following Dox and IAA treatment. (C) Scatterplots of the ratio of spindle length and cell diameter in controls (0.54 ± 0.04, n = 32) and RCC1-depleted (0.47 ± 0.04, n = 23) cells. Bars indicate means ± SDs from >3 independent experiments. * indicates statistical significance according to Welch’s *t*-test (*p* < 0.0001). (D) Quantification of cells with unfocused spindles in each condition from data in (C) and (E). Bars indicate means ± SEMs. N = 27, 34, 37, and 113 from >4 independent experiments. p-values calculated using Dunnett’s multiple comparisons test after one-way ANOVA (*F* (3,14) = 36.40, *p* < 0.0001). * indicates p < 0.0001. (E) Live fluorescent images of SiR-DNA, RCC1-mAC, NuMA-mAID-mCherry, and SiR700-TUB in RCC1-mAC and NuMA-mAID-mCh double knock-in cells following 24 hr of Dox and IAA treatment. Two cells with or without RCC1 and NuMA signals were analyzed in the same field. Eight z-section images were acquired using 1.0-μm spacing. Maximum intensity projection images are shown. (F, G) Live fluorescent images of RCC1-mAC and NuMA-mCh in RCC1-positive control (F) and RCC1-negative cells (G) treated with nocodazole and IAA, as described in Fig. S2I. * indicates RCC1-undegraded cells. (H) Scatterplots of the ratio of spindle length and cell diameter in control (0.57 ± 0.05, n = 35) and RCC1-depleted (0.50 ± 0.06, n = 30) cells. Bars indicate means ± SDs from >3 independent experiments. * indicates statistical significance according to Welch’s *t*-test (*p* < 0.0001). (I) Quantification of spindle orientation on the x-z plane in control (n = 43) and RCC1-depleted (n = 42) cells from 2 independent experiments. See Methods for the definition of parallel and tilted orientations. * indicates statistical significance according to Z-test (significance level 0.1). Scale bars = 10 μm.

### Mitotic degradation of RCC1 does not affect localization and function of NuMA at spindle poles

NuMA is transported into the nucleus during interphase (Fig. 1B) [32, 40], where it is likely released from importins by nuclear Ran-GTP. Because we found that NuMA is maintained in the nucleus following RCC1 degradation in interphase (Fig. S2E, t = - 0:10), the majority of NuMA may already have been liberated from importins by RCC1 before its degradation and may have been maintained in an active form in the nucleus, thereby producing no aberrant phenotypes in the subsequent mitosis in RCC1-depleted cells. To exclude this possibility, we next depleted RCC1 in nocodazole-arrested cells and analyzed the behavior of NuMA following nocodazole washout (For procedure, see Fig. S2I).

In RCC1-positive control cells, NuMA diffused into the cytoplasm during nocodazole arrest (Fig. 2F, t = −90), but rapidly accumulated near chromosome masses following nocodazole washout (Fig. 2F, t = 10). NuMA localized at the poles of metaphase spindles within 60 min (Fig. 2F, t = 60) and entered the nucleus following mitotic exit (Fig. 2F, t = 85). Importantly, NuMA accumulated similarly at focused spindle poles, even if RCC1 was degraded during nocodazole arrest. RCC1-mAC signals were initially detectable on chromosome masses during nocodazole-arrest (Fig. 2G, t = −90, arrow), but were reduced to undetectable levels after addition of IAA (Fig. 2G, t = 0). After nocodazole-washout, NuMA localized to focused spindle poles after ~60 min (Fig. 2G, t = 55), as observed in control cells. Cells entered anaphase with timing similar to that of control cells (Fig. S2J), but NuMA was not recruited to the nucleus after mitotic exit (Fig. 2G, t = 80).

As observed when RCC1 was degraded in asynchronous culture (Fig. 2C), the metaphase spindle became shorter when RCC1 was depleted during nocodazole-arrest (Fig. 2H). In addition, the metaphase spindle was not properly oriented to the attached culture dishes (Fig. 2I) Taken together, these results indicate that RCC1 participates in some fashion in spindle assembly in human mitotic cells, but is dispensable for NuMA localization and function at spindle poles, even if RCC1 is degraded during mitosis.

### NuMA localized at spindle poles is released from importins independently of Ran-GTP

Our results suggest that NuMA is liberated from importins in the absence of Ran-GTP (Fig. 3A). To confirm this, we next analyzed importin localization. Because importin-α has several isoforms in human cells [41], we first examined localization of endogenous importin-β in living cells by fusing it with mCherry (Fig. S3A). Unexpectedly, importin-β-mCh accumulated on kinetochore-microtubules (k-fibers) near chromosomes, but not at metaphase spindle poles (Fig. 3B top). Although this is inconsistent with the reported spindle-pole localization of importin-β [42], this result was confirmed by other visualization methods using a mAC tag and anti-importin-β antibodies (Fig. S3B,C). Importantly, RCC1 depletion diminished importin-β from k-fibers, but did not cause importin-β accumulation at the spindle poles (Fig. 3B, bottom) where NuMA localized (Fig. 2B). This suggests that NuMA is released from importin-β at the spindle poles, even in the absence of Ran-GTP (Fig. 3A).

**Figure 3.**
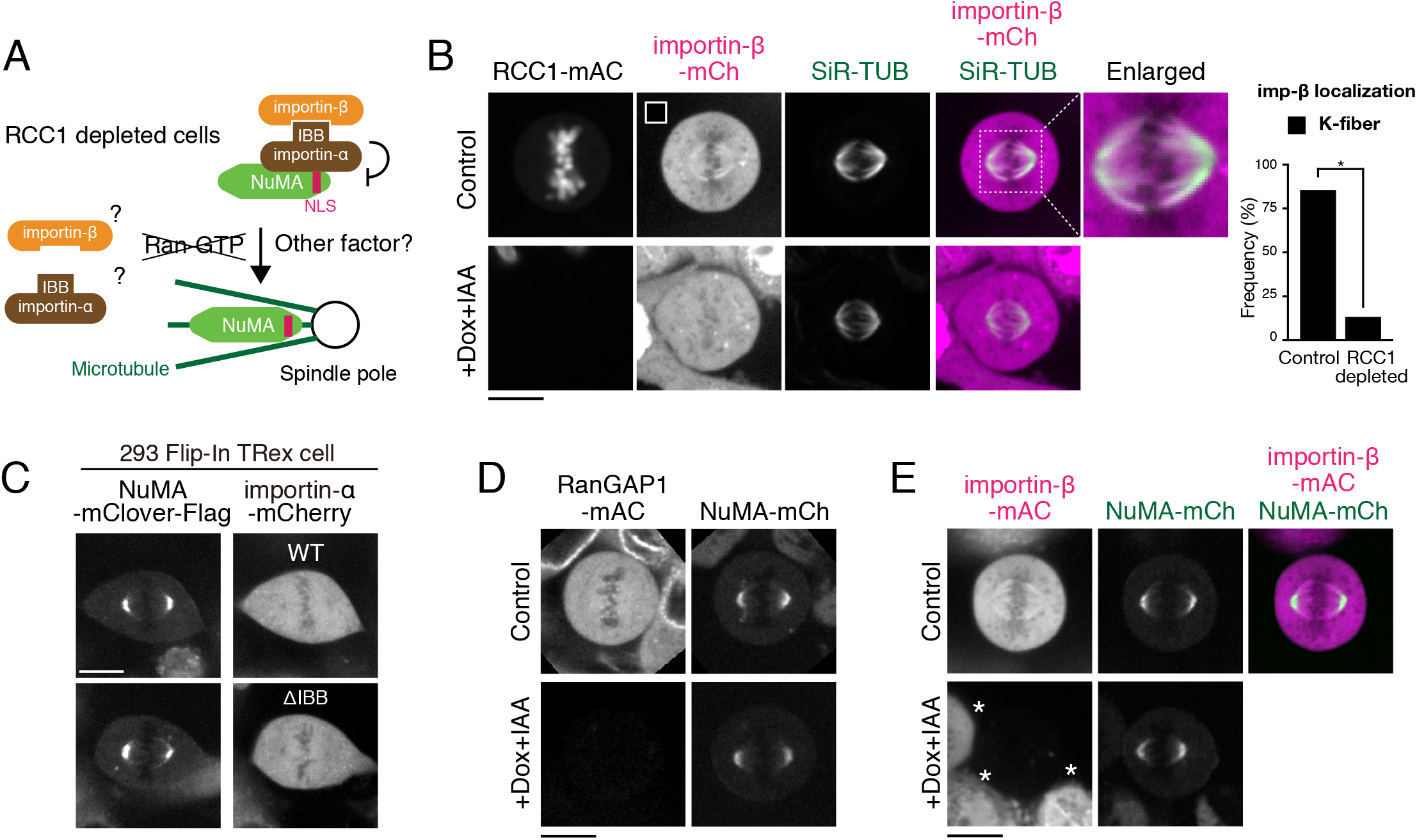
NuMA is liberated from importins at spindle poles independently of Ran-GTP. (A) A model showing NuMA liberation from importins in RCC1 depleted cells. (B) Metaphase RCC1-mAC cells showing live fluorescent images of RCC1-mAC, importin-β-mCh, and SiR-TUB and after 24 hr following treatment with Dox and IAA. Right: Quantification of cells with k-fiber localization of importin-β in control (n > 40) and RCC1-depleted (n > 40) cells from 3 independent experiments. * indicates statistical significance according to Z-test (significance level 0.0001). (C) Live fluorescent images of NuMA-mClover-FLAG (mCF, left) and importin-α (right) wild type (WT, top) and a ΔIBB mutant (bottom). (D-E) Metaphase RanGAP1-mAC (D) and importin-β-mAC (E) cells showing live fluorescent images of NuMA-mCherry (mCh), SiR-tubulin (SiR-TUB), and RanGAP1-mAC (D) or importin-β-mAC (E) after 24 hr following treatment with Dox and IAA. * in E indicates cells with importin-β signals in the presence of Dox and IAA. Scale bars = 10 μm.

To further test whether Ran-independent pathways exist for NuMA activation, we next analyzed localizations of importin-α wild type (WT) and ΔIBB mutants, which lack the importin-β-binding (IBB) domain. Importin-α ΔIBB mutants are insensitive to Ran-GTP due to the lack of an IBB domain, but are still able to interact with NuMA and partially inhibit NuMA’s microtubule-binding activity *in vitro* (Fig. 3A) [32]. However, importin-α ΔIBB diffused into cytoplasm similarly to importin-α WT, and neither affected NuMA’s spindle-pole localization nor colocalized with NuMA at the spindle poles in our experimental conditions (Fig. 3C, S3D). These results suggest that NuMA is released from the importin-α/β complex in a Ran-GTP-independent manner and that it localizes at spindle poles.

### Ran-GAP1 and importin-β degradation do not affect NuMA localization and function at spindle poles

Although RCC1 depletion does not affect NuMA localization and functions, degradation of Ran-GAP1 or importin-β may cause abnormal activation of NuMA throughout human cells, resulting in spindle assembly defects. To test this, we next degraded either Ran-GAP1 or importin-β using AID technology (Fig. 3D-E). Ran-GAP1 degradation caused few mitotic phenotypes (Fig. 3D, S3E-I) and did not affect NuMA’s spindle-pole localization (Fig. 3D bottom). Similarly, importin-β degradation did not affect NuMA’s localization and function at spindle poles (Fig. 3E bottom), although importin-β degradation caused short spindles and mitotic delay (Fig. S3J-N). These results indicate that Ran-dependent spatial regulation is dispensable for NuMA localization and function at spindle poles in cultured human cells.

### RCC1 regulates chromosome-proximal localization of HURP and HSET

RCC1 depletion caused shorter mitotic spindles (Fig. 2B, C, F, H), suggesting that Ran-GTP serves some function in spindle assembly in human cells. To identify spindle assembly factors (SAFs) regulated by Ran-GTP, we next analyzed the localization of 3 other major SAFs: TPX2, HSET, and HURP. mCherry-tagged TPX2 colocalized with SiR-tubulin signals in metaphase (Fig. 4A top, Fig. S4A), and its localization was virtually unaffected in RCC1-depleted cells (Fig. 4A bottom), as observed for NuMA (Fig. 2B). In contrast, mCherry-tagged HSET localized everywhere along spindle microtubules (Fig. 4B top, Fig. S4B) [30], and its spindle localization was selectively reduced near chromosomes following RCC1 depletion, although HSET still localized along spindle fibers farther away from chromosomes (Fig. 4B bottom). On the other hand, mCherry-tagged HURP accumulated at k-fibers near chromosomes, but localized weakly on spindle microtubules following RCC1 depletion (Fig. 4C and Fig. S4C). These results suggest that in human mitotic cells, the chromosome-derived Ran-GTP gradient regulates SAF localization preferentially near chromosomes, regardless of the presence of NLS (Fig. 4D).

**Figure 4.**
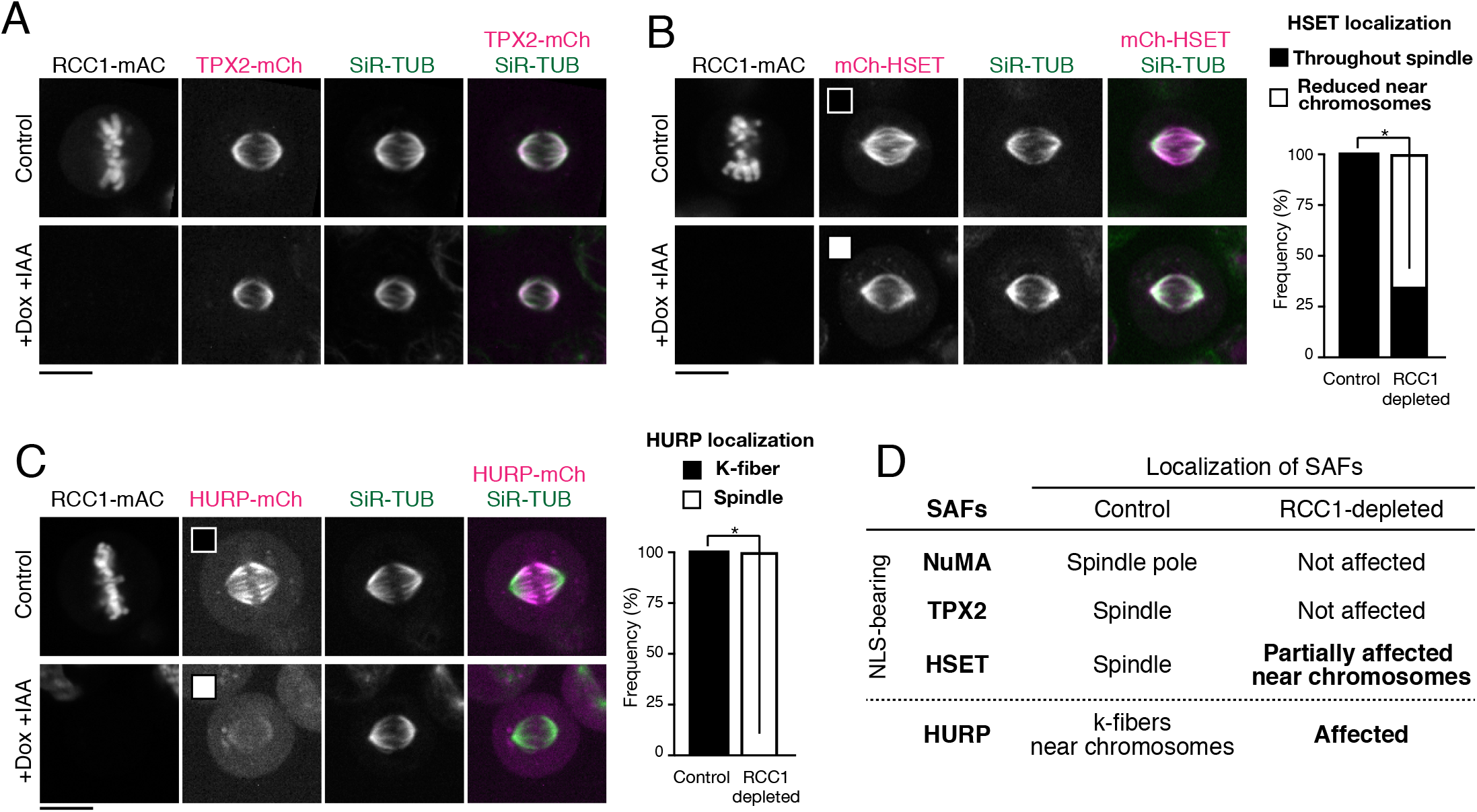
RCC1 regulates chromosome-proximal localization of HURP and HSET. (A-C) Left: Metaphase RCC1-mAC cells showing live fluorescent images of RCC1-mAC, SiR-TUB and TPX2-mCh (A), mCh-HSET (B), and HURP-mCh (C) after 24 hr following treatment with Dox and IAA. Right: Quantification of throughout spindle or k-fiber localization of HSET or HURP in control (n > 30) and RCC1-depleted (n > 40) cells from 2 or 3 independent experiments. * indicates statistical significance according to Z-test (significance level 0.0001). (D) A list summarizing localization of SAFs in control and RCC1-depleted cells. Scale bars = 10 μm.

### HURP, but not importin-β, is required to stabilize k-fibers

Importin-β inhibits HURP’s microtubule-binding activities by masking one of HURP’s microtubule-binding domains (MTBD2) [43] (Fig. 5J). To understand the relationship between HURP and importin-β for k-fiber localization and function, we next sought to degrade endogenous HURP using AID (Fig. 5A and Fig. S5A-B). Endogenous HURP-mACF accumulated at k-fibers near chromosomes (Fig. 5A top), as observed with anti-HURP antibodies [31]. HURP depletion resulted in diminished importin-β localization to k-fibers (Fig. 5A-B, S5C) and reduced mitotic spindle length (Fig. 5C). Because k-fibers are resistant to cold treatment [31], we next incubated cells with ice-cold medium for 20 min and analyzed cold-stable microtubules. HURP localized to cold-stable microtubules (Fig. 5D, top), which were disrupted by HURP depletion (Fig. 4D bottom), consistent with a previous study [31].

**Figure 5.**
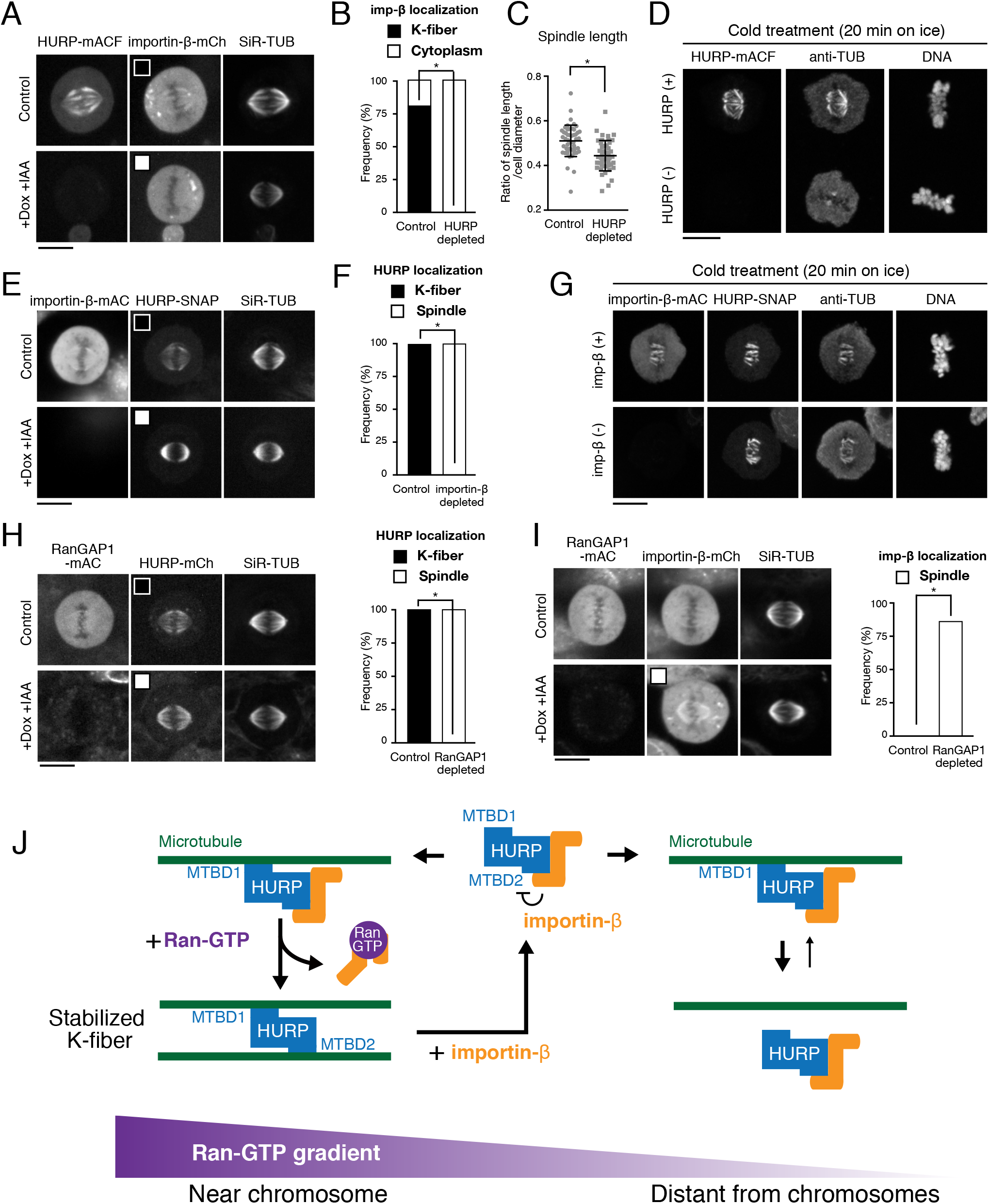
HURP, but not importin-β, is required to stabilize k-fibers. (A) Metaphase HURP-mACF cells showing live fluorescent images of HURP-mACF, importin-β-mCh and SiR-TUB after 24 hr following Dox and IAA treatment. (B) Quantification of k-fiber localization of importin-β in control (n = 49) and HURP-depleted (n = 46) cells from 3 independent experiments. (C) Scatterplots of the ratio of spindle length and cell diameter in control (0.64 ± 0.05, n = 49) and HURP-depleted (0.52 ± 0.06, n = 43) cells. * indicates statistical significance according to Welch’s *t*-test (*p* < 0.0001). (D) Fluorescent images of HURP-mACF, TUB, and DNA (Hoechst 33342 staining) in metaphase fixed cells treated with ice-cold medium for 20 min. Two cells with or without HURP signals were analyzed in the same field. (E) Metaphase importin-β-mAC cells showing live fluorescent images of importin-β-mAC, HURP-SNAP and SiR-TUB after 24 hr following treatment with Dox and IAA. (F) Quantification of spindle localization of HURP in control (n = 49) and importin-β-depleted (n = 43) cells from 3 independent experiments. (G) Fluorescent images of importin-β-mAC, HURP-SNAP, TUB, and DNA (Hoechst 33342 staining) in metaphase fixed cells treated with ice-cold medium for 20 min. Five z-section images were obtained using 0.5-μm spacing and maximum intensity projection images are shown in (D) and (G). (H-I) Left: metaphase RanGAP1-mAC cells showing live fluorescent images of RanGAP1-mAC, SiR-TUB and HURP-mCh (H) or importin-β-mCh (I) after 24 hr following Dox and IAA treatment. Right: quantification of k-fiber localization of HURP or importin-β in control (n = 45) and RanGAP1-depleted (n > 45) cells from 3 independent experiments. * in (B), (F), (H) and (I) indicates statistical significance according to Z-test (significance level 0.0001). (J) A local cycling model of HURP on k-fibers regulated by Ran-GTP and importin-β. See text for details. Scale bars = 10 μm.

We next depleted importin-β and analyzed effects of this depletion on HURP and k-fibers (Fig. 5E, S5D). Importin-β depletion caused a remarkable re-localization of HURP from k-fibers near chromosomes to spindle microtubules (Fig. 5E-F). Although k-fiber localization of HURP was unclear in importin-β-depleted cells due to the relatively strong accumulation of HURP on spindle microtubules around spindle poles (Fig. 5E bottom), HURP was clearly detected on cold-stable k-fibers in importin-β-depleted cells (Fig. 5G bottom). These results suggest that HURP acts in k-fiber stabilization, independently of importin-β.

### HURP and importin-β localize throughout the spindle in RanGAP1-depleted cells

Whereas HURP and importin-β have different roles in k-fiber stabilization (Fig. 5D, G), both proteins accumulate at k-fibers near chromosomes downstream of RCC1 (Fig. 3B, 4C). To better understand mechanisms of Ran-based spatial regulation of HURP and importin-β, we next analyzed behavior of HURP and importin-β in RanGAP1-depleted cells, in which Ran-GTP should exist throughout cells. Interestingly, both HURP and importin-β localized throughout the spindle with increased intensities in RanGAP1-depleted cells (Fig. 5H-I, S5E). These results suggest that HURP and importin-β act together and interact with microtubules preferentially in the presence of Ran-GTP (Fig. 5J).

### HURP dynamically associates with k-fibers in the presence of importin-β

Based on our results, we developed a local cycling model for activation and polarization of HURP (Fig. 5J). In this model, importin-β inhibits HURP globally, including at k-fibers, by masking HURP’s 2^nd^ microtubule-binding domain (MTBD2). The resulting HURP-importin-β complex binds weakly to microtubules through HURP’s MTBD1 [43], but the Ran-GTP gradient locally releases importin-β from HURP, resulting in full activation of HURP near chromosomes (Fig. 5J). To test this model, we first performed fluorescence recovery after photobleaching (FRAP) for HURP, and analyzed its dynamics on spindle microtubules in the presence and absence of importin-β. In control cells, HURP was quickly recovered at k-fibers after bleaching (Fig. 6A top, 6B black, S6A t½ = 20.5 sec). In contrast, HURP’s fluorescent signals were hardly seen on the spindle in importin-β-depleted cells (Fig. 6A bottom, 6B red, S6B). These results indicate that HURP dynamically associates with k-fibers in the presence of importin-β, whereas HURP binds tightly to spindle microtubules in the absence of importin-β.

**Figure 6.**
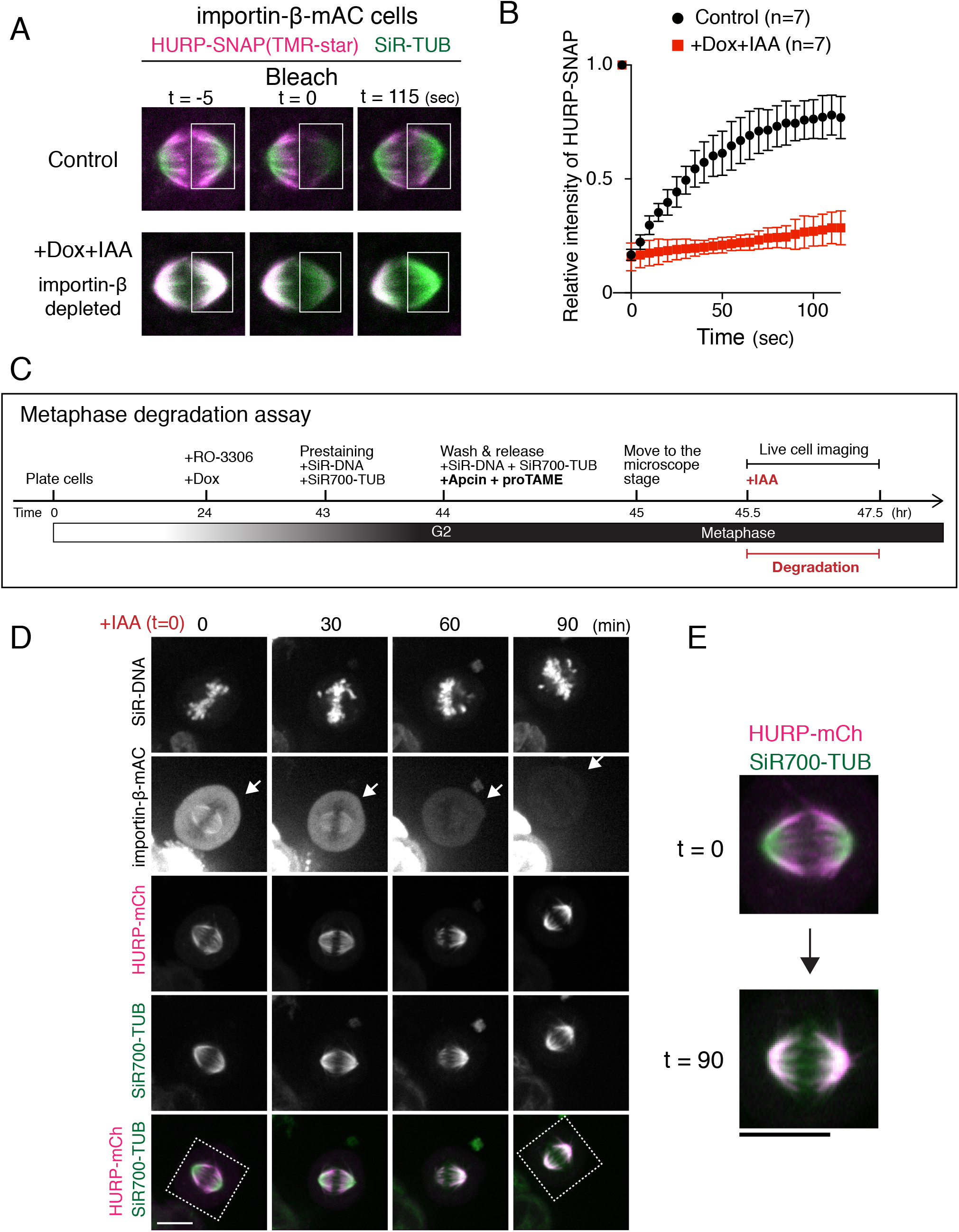
HURP dynamically accumulates on metaphase k-fibers in an importin-β-dependent manner. (A) Live fluorescent images of HURP-SNAP visualized with TMR-star (magenta) and SiR-tubulin (TUB) in control (top) and importin-β-depleted cells (bottom). Fluorescent signals were bleached in the indicated box region at t = 0, and the fluorescence recoveries were monitored for 120 sec. (B) A graph showing fluorescence recovery after photobleaching. An average of 7 samples was plotted. Bars indicate SDs. (C) Schematic diagram of the metaphase degradation assay. Following release from RO-3336-mediated G2 arrest, proTAME and Apcin were added to arrest cells in metaphase. Auxin (IAA) was added (indicated by the red line) to induce RCC1 degradation during metaphase. (D) Live fluorescent images of SiR-DNA, importin-β-mAC, HURP-mCh, and SiR-700-tubulin (TUB). IAA was added at t = 0. Arrows indicate a cell showing a reduction of importin-β signal during metaphase. (E) Enlarged images from (D) showing a re-localization of HURP-mCh from k-fibers (t = 0) to the spindle (t =90). Scale bars = 10 μm.

### HURP is dynamically maintained at k-fibers during metaphase

mAID-tag fusion proteins can be rapidly degraded with a half-life of 20 min [35]. To confirm the dynamic regulation of HURP by importin-β and Ran-GTP, we next sought to degrade importin-β during metaphase by combining AID-mediated degradation with APC/C inhibitors [44] (Fig. 6C). Following treatment with the APC/C inhibitors, Apcin and proTAME, cells arrested at metaphase, in which both importin-β and HURP accumulated at k-fibers near chromosomes (Fig. 6 D, t = 0). Importantly, importin-β-mAC signals diminished to undetectable levels 60-90 min after addition of IAA (Fig. 6D, arrows), and HURP relocated from k-fibers to spindle microtubules in response to the reduction of importin-β signals (Fig. 6D, E).

To confirm these results, we next acutely degraded RCC1 in metaphase-arrested cells. As with importin-β degradation, HURP dissociated from k-fibers and localized weakly on the spindle in response to degradation of RCC1 (Fig. 7A, B). Unexpectedly, in contrast to the prometaphase degradation assay (Fig. 2H), spindle length appeared normal when RCC1 was degraded in the metaphase-arrested condition (Fig. 7C). Together, these results indicate that HURP is dynamically maintained at k-fibers near chromosomes by the Ran-Importin pathway, even in metaphase, but HURP is critical for spindle length regulation primarily during prometaphase.

**Figure 7.**
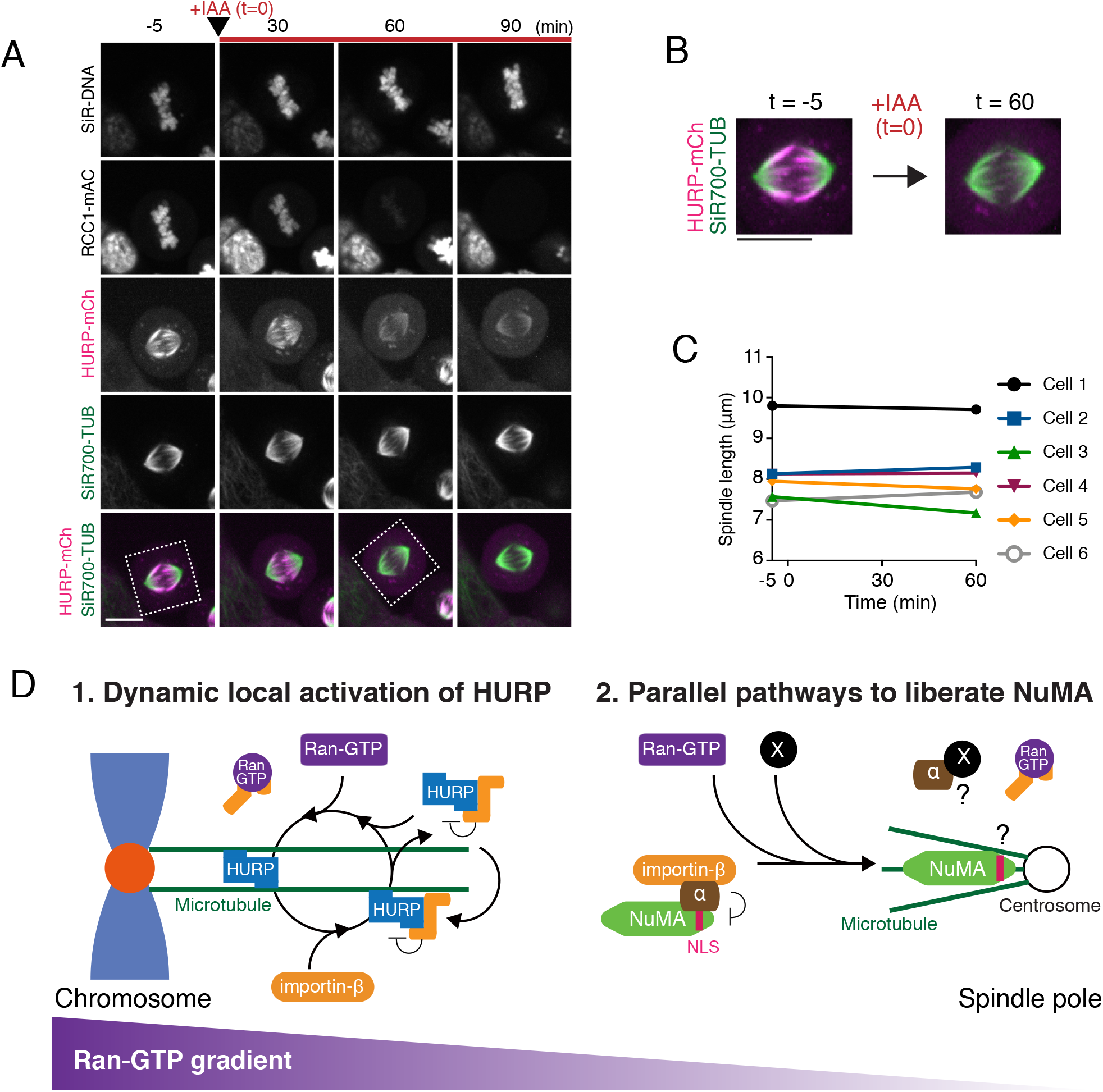
Models of local activation mechanisms for HURP and NuMA in mitosis. (A) Live fluorescent images of SiR-DNA, RCC1-mAC, HURP-mCh, and SiR-700-tubulin (TUB). IAA was added at t = 0. (B) Enlarged images of indicated regions in (A) showing a reduction of HURP-mCh from k-fibers in response to degradation of RCC1. (C) Spindle length measurement (n = 6) at t = −5 and 60 min in (A). (D) Left: in the vicinity of chromosomes, Ran-GTP and importin-β promote the microtubule binding and dissociation cycle of HURP, resulting in polarized HURP accumulation and stable k-fiber formation. Right: chromosome-derived Ran-GTP is not required to activate NuMA at the spindle poles in mitotic human cells. A Ran-independent, parallel pathway would exist to activate NuMA away from chromosomes. See text for details. Scale bars = 10 μm.

## Discussion

### NuMA is liberated from importins independently of Ran-GTP for spindle-pole focusing in human mitotic cells

In contrast to the prevailing model (Fig. 2A), we demonstrated that the Ran-Importin pathway is dispensable for localization and functions of NuMA at the spindle poles in human HCT116 cells (Fig. 2, 3, 7D right). This is consistent with the recent observation that NuMA is less sensitive to Ran-GTP level than to HSET/XCTK2 [29]. Although we do not exclude the possibility that Ran-GTP liberates NuMA from importin-α/β complexes near chromosomes, we favor the idea that parallel pathways exist to activate NuMA in mitotic human cells. In fact, recent studies indicate that importin-α/β-binding TPX2 can be activated not only by Ran-GTP, but also by Golgi-or palmitoylation-dependent sequestration of importin-α [45, 46]. In addition, mitotic spindles contain centrosomes, which may generate special signals that liberate NuMA from inhibitory importins (Fig. 7D). Interestingly, NuMA is broadly distributed on a bundle-like structure between the poles in human acentrosomal cells [47]. It is necessary to analyze whether NuMA is preferentially regulated by Ran-GTP in acentrosomal cells, especially in oocytes, where Ran-GTP governs meiotic spindle assembly [12].

Although NLS-containing SAFs are recognized by importin-α, structural studies indicate that importin-α binds to NuMA and TPX2 with slightly different binding patches [32]. In addition, whereas TPX2-NLS and NLS-binding sites of importin-α are well conserved in vertebrates, NLS of NuMA is not well conserved in fish (Fig. S1D-F). Furthermore, NLS is lacking in other NuMA-like proteins in lower eukaryotes, such as *Caenorhabditis elegans* LIN-5, *Drosophila* Mud, and yeast Num1 [38, 48, 49], suggesting that NuMA acquired NLS in higher animals and is likely to be regulated differently than TPX2. Future research should be undertaken to understand how the NuMA-importin interaction is regulated in a Ran-independent manner, and why NLS-dependent regulation of NuMA was acquired in higher animals.

### The Ran-Importin pathway locally activates and polarizes HURP by promoting its microtubule binding-dissociation cycle near chromosomes

In contrast to NuMA, we demonstrated that HURP is preferentially regulated by the Ran-Importin pathway in mitotic human cells (Fig. 4C, 5E, H). Although HURP has been identified previously as a downstream target of Ran-GTP [31], we unexpectedly found that HURP also colocalizes with importin-β on k-fibers near chromosomes (Fig. S3C, Fig. 5A, E), and stabilizes k-fibers independently of importin-β (Fig. 5D, G). In addition, HURP’s spindle distribution is sensitive to levels of Ran-GTP and importin-β (Fig. 4C, 5E, H), and is dynamically and spatially maintained during metaphase in a Ran-pathway-dependent manner (Fig. 6A, B, D, 7A). Based on these results, we propose a local cycling model for establishment and maintenance of HURP’s polarized localization to spindle microtubules (Fig. 5J, 7D left). After nuclear envelope breakdown (NEBD), HURP strongly interacts with microtubules through its two microtubule-binding domains (MTBD1 and MTBD2 in Fig. 5J) [31, 43]. Since importin-β is localized diffusely throughout cells (Fig. 3E, 5E), it binds to HURP on microtubules, and then dissociates HURP from the microtubules by masking HURP’s MTBD2 domain [43]. However, in the vicinity of chromosomes, Ran-GTP releases HURP from importin-β [31], and the liberated HURP interacts strongly with microtubules around chromosomes. By repeating this local binding-dissociation cycle, HURP, but not importin-β, stabilizes microtubules and generates stable k-fibers near chromosomes (Fig. 5D, G). This dynamic regulation is similar to that of HSET/XCTK2 [50] and would be suitable for bundling short microtubules around kinetochores during prometaphase [51] and for coupling HURP’s polarized localization with microtubule flux on the metaphase spindle.

### RCC1 is required to define proper spindle length during prometaphase

RCC1 depletion causes shortened bipolar spindles in human cells (Fig. 2B, C, F, H). This is probably due to multiple defects in spindle assembly processes, including the lack of HURP-based k-fiber formation (Fig. 5C, D) and HSET-dependent microtubule-sliding (Fig. 4B) [28]. Intriguingly, our mitotic degradation assays indicate that Ran-GTP controls spindle length primarily during prometaphase, rather than in metaphase (Fig. 2H, 7C). Once metaphase spindles are assembled, other k-fiber localized proteins, such as clathrin, TACC3, and ch-TOG [52], may be able to maintain bundled-k-fibers during metaphase in a Ran-independent manner.

In addition, our mitotic degradation assay revealed that RCC1-depletion does not affect mitotic progression (Fig. S2J). This suggests that the mitotic delay observed in RCC1 depletion in asynchronous culture (Fig. S2F) is a secondary defect caused by loss of interphase RCC1 activity. In fact, ectopically expressed HSET-NLS mutants localize in cytoplasm and causes abnormal cytoplasmic microtubule-bundling in interphase [30]. Numerous similar defects would be created by RCC1 depletion in interphase and would affect subsequent mitotic progression.

### *A* new toolkit and mitosis-specific degradation assays to dissect mitotic roles of the Ran-importin pathway

As discussed above, mitotic inactivation is critical to precisely analyze mitotic functions of Ran-GTP and importins. Previously, tsBN2, a temperature-sensitive RCC1 mutant hamster cell line [53, 54] and a small molecule inhibitor, importazole [55], have been developed to acutely inhibit functions of RCC1 and importin-β, respectively. Here, we established three human AID-cell lines for RCC1, RanGAP1, and importin-β [35], and succeeded in degrading RCC1 and importin-β specifically in prometaphase (Fig. 2C-F) or metaphase (Fig. 6C-E, 7A-B). Because these AID-cell lines and mitotic degradation assays are applicable to other Ran-regulated SAFs/cortical proteins [4, 54, 56] and other multi-functional proteins such as dynein and NuMA [20, 35], respectively, these toolkits and assays will further advance our understanding of mechanisms and roles of spindle assembly, maintenance, and positioning in animal cells.

## Acknowledgments

We thank Iain M. Cheeseman for critical reading of the manuscript, and Yuki Tsukada, Rie Inaba and Kiyoko Murase for technical assistance. This work was supported by grants from the PRESTO program (JPMJPR13A3) of the Japan Science and Technology agency (JST) for T.K, a Career Development Award of the Human Frontier Science Program (CDA00057/2014-C) for T.K., KAKENHI (16K14721 and 17H05002 for T.K, 17H01431 for G.G.) of the Japan Society for Promotion of Science (JSPS), NIG-JOINT (2014B-B-3, 2015-A1-19, 2016-A1-22 for T.K.) of National Institute of Genetics (NIG), the Naito Foundation for T.K, and JSPS and DFG under the Joint Research Projects-LEAD with UKRI for G.G.

## Author contributions

Conceptualization, TK; Investigation, TK, KT, HH, MN, and MO; Formal analysis, TK and KT; Methodology, TK, YS, and MK; Writing, TK; Supervision, TK and GG; Funding Acquisition, TK and GG.

## Declaration of interests

The authors declare no competing interests.

## Materials and Methods

- Plasmid Construction Plasmids for CRISPR/Cas9-mediated genome editing and auxin-inducible degron were constructed according to protocols of Natsume et al. [35] and Okumura et al., [20]. To construct donor plasmids containing homology arms for RCC1 (~500-bp homology arms), RanGAP1 (~500-bp), importin-β (~500-bp), HURP (~200-bp), TPX2 (~200-bp), and HSET (~200-bp), gene synthesis services from Eurofins Genomics K.K. (Tokyo, Japan) or Genewiz (South Plainsfield, NJ) were used. Plasmids and sgRNA sequences used in this study are listed in Supplementary Tables S1 and S2, and will be deposited in Addgene.
- Cell Culture, Cell Line Generation, and Antibodies HCT116 cells were cultured as described previously [20]. Knock-in cell lines were generated according to procedures described in Okumura et al. [20]. To activate auxin-inducible degradation, cells were treated with 2 μg/mL Dox and 500 μM indoleacetic acid (IAA) for 20–24 hr. Cells with undetectable signals for mAID-fusion proteins were analyzed. Flip-In T-REx 293 cells were used in Figure 3C to express mCherry-tagged importin-α constructs. Cell lines were created according to procedures described in Kiyomitsu et al. [57]. To induce transgenes, cells were incubated with 1 μg/mL tetracycline (MP Biomedicals). Cell lines and primers used in this study are listed in Tables S1 and S3, respectively. Antibodies against tubulin (DM1A, Sigma-Aldrich, 1:2,000), NuMA (Abcam, 1:1,000), RCC1 (Cell Signaling Technology, D15H6, Rabbit mAb, 1:100), RanGAP1 (Santa Cruz Biotechnology, H-180, 1:200), importin-β (GeneTex, 3E9 Mouse mAb, 1:100), and HURP (E. Nigg laboratory, 1: 200) were used for western blotting. For RCC1 immunoblots, membranes were incubated with anti-RCC1 antibody overnight at 4 °C.
- Microscope System Imaging was performed using spinning-disc confocal microscopy with a 60× 1.40 numerical aperture objective lens (Plan Apo λ, Nikon, Tokyo, Japan). A CSU-W1 confocal unit (Yokogawa Electric Corporation, Tokyo, Japan) with five lasers (405, 488, 561, 640, and 685 nm, Coherent, Santa Clara, CA) and an ORCA-Flash 4.0 digital CMOS camera (Hamamatsu Photonics, Hamamatsu City, Japan) were attached to an ECLIPSE Ti-E inverted microscope (Nikon) with a perfect focus system. DNA images in Figure 2A/B or Figure 4D/G were obtained using either a SOLA LED light engine (Lumencor, Beaverton, OR) or a 405-nm laser, respectively.
- Immunofluorescence and Live Cell Imaging For immunofluorescence in Figure S1K, HURP-mACF cells were fixed with PBS containing 3% paraformaldehyde and 2% sucrose for 10 min at room temperature. Fixed cells were permeabilized with 0.5% Triton X-100™ for 5 min on ice, and pretreated with PBS containing 1% BSA for 10 min at room temperature after washing with PBS. Importin-β was visualized using anti-importin-β antibody (1:500). Images of multiple z-sections were acquired by spinning-disc confocal microscopy using 0.5-μm spacing and camera binning 2. Maximally projected images from 3 z-sections are shown. For live cell imaging, cells were cultured on glass-bottomed dishes (CELLview™, #627860 or #627870, Greiner Bio-One, Kremsmünster, Austria) and maintained in a stage-top incubator (Tokai Hit, Fujinomiya, Japan) to maintain the same conditions used for cell culture (37° C and 5% CO2). In most cases, three to five z-section images using 0.5-μm spacing were acquired and single z-section images are shown, unless otherwise specified. Microtubules were stained with 50 nM SiR-tubulin or SiR700-tubulin (Spirochrome) for >1 hr prior to image acquisition. DNA was stained with 50 ng/mL Hoechst^®^ 33342 (Sigma-Aldrich) or 20 nM SiR-DNA (Spirochrome) for > 1 hr before observation. To visualize SNAP-tagged HURP in Fig. 4E, cells were incubated with 0.1 μM TMR-Star (New England BioLabs) for > 2 hr, and TMR-Star were removed before observation. To optimize image brightness, the same linear adjustments were applied using Fiji and Photoshop.
- Prometaphase degradation assay and nocodazole washout To degrade mAID-tagged proteins during nocodazole arrest, cells were treated with 2 μg/mL Dox and 3.3 μM nocodazole at the indicated times (Fig. 2C). Five hours after addition of nocodazole, cell culture dishes were moved to the stage of a microscope equipped with a peristaltic pump (SMP-21S, EYELA, Tokyo Rikakikai). Two z-section images were acquired using 2-μm spacing at three different (X.Y) positions and at 5-min intervals, with 500 μM IAA added during the first interval. After 90 min, the nocodazole-containing medium was completely replaced with fresh medium using the peristaltic pump at a velocity of 20 sec/mL for 15 min. Images were acquired for a further 2 hr and maximum intensity projection images are shown in Figure 2D-F. To analyze spindle orientation in Figure 2I, we took five z-section images using 2-μm spacing. When both spindle poles are included within three z-section images, we judged the spindle as having parallel orientation.
- Metaphase degradation assay To degrade mAID-tagged proteins in metaphase-arrested cells, cells were treated with 50 μM Apcin (I-444, Boston Biochem) and 20 μM proTAME (I-440, Boston Biochem) at the indicated times (Fig. 5A). Three z-section images were acquired using 1-μm spacing at six different (X,Y) positions and at 5-min intervals, with 500 μM IAA added during the first interval. Maximum intensity projection images are shown in Figure 5B.
- Cold treatment assay To increase the number of cells in metaphase, cells were treated with 20 μM MG132 (C2211, Sigma-Aldrich) for 90 min. To visualize SNAP-tagged HURP (Fig. 4G), cells were incubated with 0.1 μM TMR-Star (S9105S, New England BioLabs) for at least 30 min. Before fixation, cells were incubated in ice-cold medium for 20 min [31] to depolymerize non-kinetochore microtubules.
- FRAP FRAP was conducted with a microscope (LEM 780, Carl Zeiss MicroImaging, Inc.), using a 63 x objective lens. Images were acquired every 5 sec before and after photobleaching. The bleached area (BA) was set as it covers half spindle and illuminated at t = 0 using 560 nm laser (20 mW) with the following setting: speed 4.0 and iteration 1. Metaphase cells that orient parallel to the bottom cover-glass were selected. HURP (TMR-Star) intensity of BA was normalized using the intensity of non-bleached area (NBA) that covers the remaining half spindle. Corrected relative intensity at time t_n_ was calculated as (BA_n_ – BG_n_) / (BA_-1_ – BG_-1_) x (NBA_-1_ – BG_-1_) / (NBA_n_ – BG_n_), where t = −1 represents the first time point of image acquisition before bleaching. BG means background [58]. Curve fitting and analyses shown in Fig. S6 were performed using Fiji.
- Statistical Analysis To determine the significance of differences between the mean values obtained for two experimental conditions, Welch’s *t*-tests (Prism 6; GraphPad Software, La Jolla, CA) or a Z-test for proportions (epitools.ausvet.com.au/ztesttwo) were used, as indicated in figure legends.

## Supplemental Information

### Supplemental Figure Legends

**Figure S1.**
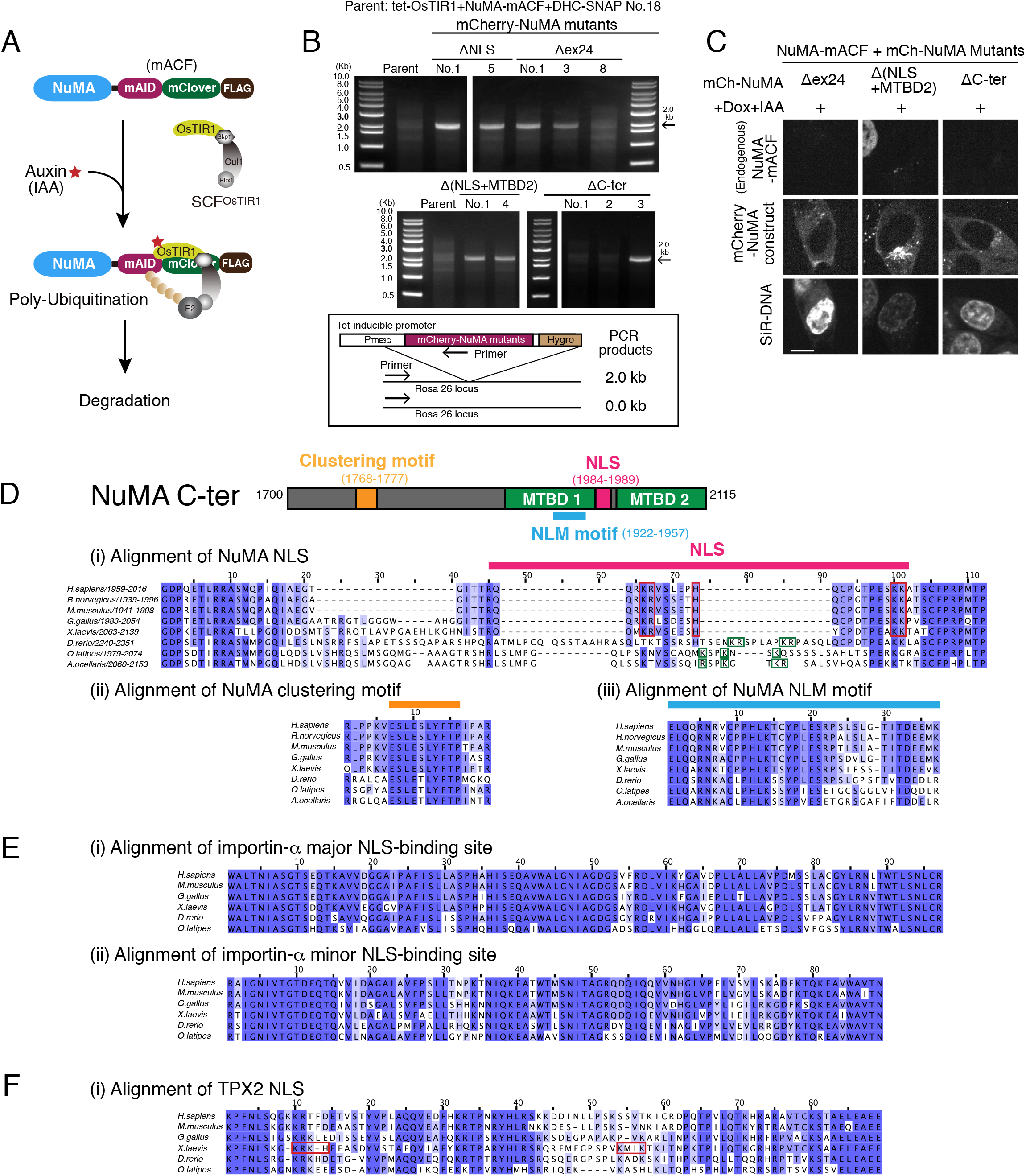
Generation of cell lines that conditionally degrade endogenous NuMA and express NuMA mutants. (A) Schematic of the auxin-inducible degradation (AID) system. OsTIR1, an F-box protein expressed following Dox treatment, forms SCF E3 ubiquitin ligase complexes. Following auxin (IAA) treatment, mAID-fusion protein was poly-ubiquitinated by SCF^OsTIR1^ and degraded by proteasomes with a half-life of ~20 min. (B) Genomic PCR showing clone genotypes after hygromycin (Hygro) selection. Clones used in this study are listed in Table S1. (C) Interphase NuMA-mACF cell lines showing live fluorescent images of NuMA-mACF, NuMA-mCh mutants, and SiR-DNA after 24 hr following treatment with Dox and IAA. Endogenous NuMA-mACF signals were undetectable, whereas ectopically expressed NuMA mutants were detected in cytoplasm. NuMA-mCh Δ(NLS+MTBD2) appeared to accumulate on microtubules around centrosomes. (D) Amino acid sequence alignment of the NLS of NuMA proteins in *H. sapiens* (NP_006176), *R. norvegicus* (NP_001094161), *M. musculus* (NP_598708), *G. gallus* (NP_001177854), *X. laevis* (NP_001081559), *D. rerio* (NP_001316910), *O. latipes* (XP_020564048), and *A. ocellaris* (XM_023273896) aligned by ClustalWS. NLSs are not well conserved in fish, although NuMA clustering motif (shown in orange) and NLM motif (sky blue) are highly conserved in vertebrates. In the NLS alignment, key amino acids that interact with importin-α [32] are boxed in red, whereas positively charged amino acids in fish are boxed in green. (E) Amino acid sequence alignment of the major (i) and minor (ii) NLS-binding site of importin-α proteins in *H. sapiens* (NP_001307540), *M. musculus* (NP_034785), *G. gallus* (NP_001006209), *X. laevis* (NP_001080459), *D. rerio* (NP_001002335), and *O. latipes* (XP_023816136) aligned by ClustalWS. (F) Amino acid sequence alignment of the NLS of TPX2 proteins in *H. sapiens* (NP_036244), *M. musculus* (NP_001135447), *G. gallus* (NP_989768), *X. laevis* (AAH68637), *D. rerio* (NP_001314674), and *O. latipes* (XP_020557297) aligned by ClustalWS. Key amino acids interact with importin-α [33] are boxed in red.

**Figure S2.**
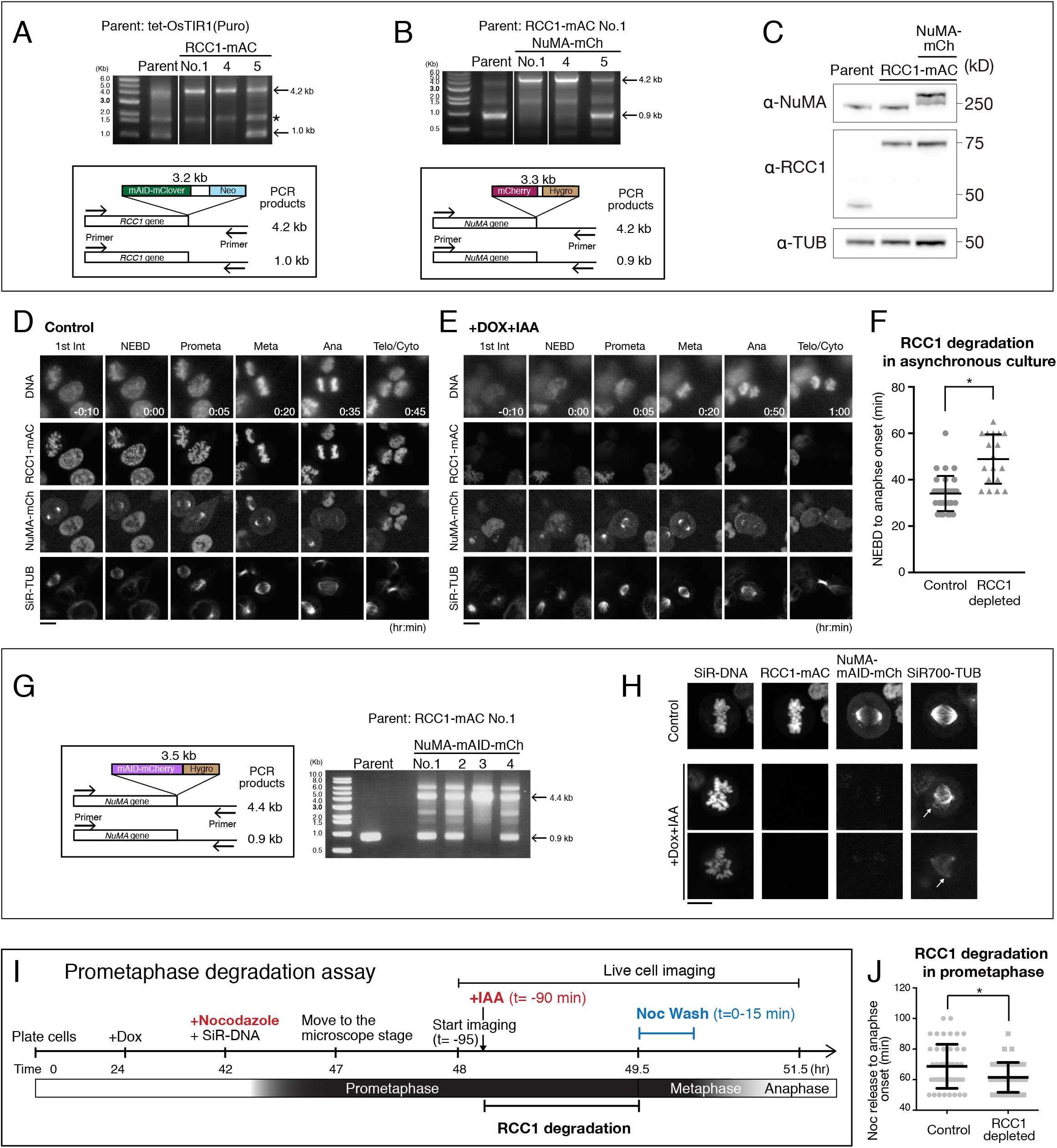
Generation of cell lines for auxin-inducible degradation of endogenous RCC1. (A) Genomic PCR showing clone genotypes after neomycin (Neo) selection. Clone No.1 was used as a parental cell in subsequent selections. * indicates a non-specific band. (B) Genomic PCR showing clone genotypes after hygromycin (Hygro) selection. Clone No.1 was used in this study. (C) Immunoblotting for anti-NuMA, anti-RCC1, and anti-α-tubulin (TUB, loading control) showing bi-allelic insertion of the indicated tags. (D-E) Live fluorescent images of DNA (Hoechst 33342 staining), RCC1-mAC, NuMA-mCh, and SiR-TUB in control (D) and RCC1-depleted (E) cells. (F) Scatterplots of mitotic duration (NEBD to anaphase onset) in control (34.1 ± 7.6, n=32) and RCC1-depleted cells (47.2 ± 10.5, n=27). Bars indicate means ± SDs from >3 independent experiments. * indicates statistical significance according to Welch’s *t*-test (p<0.0001). (G) Genomic PCR showing clone genotype after hygromycin (Hygro) selection. Clone No.3 was selected for further use. (H) Live fluorescent images of SiR-DNA, RCC1-mAC, NuMA-mAID-mCh, and SiR-TUB. A spindle-pole focusing defect (indicated by the arrow in panel 2) and abnormal spindle formation (panel 3) were observed in RCC1-mAC and NuMA-mAID-mCh co-depleted cells 20-24 hr after Dox and IAA treatment. Five z-section images were acquired using 1.0-μm spacing and maximum intensity projection images are shown. (I) (C) Schematic diagram of the prometaphase degradation assay. Nocodazole was added to arrest the cells in prometaphase, and then Auxin (IAA) was added to induce RCC1 degradation during nocodazole-arrest. Nocodazole were washed out by changing medium for 15 min with peristaltic pumps, while recording the cells. See Methods for details. (J) Scatterplots of mitotic duration (from nocodazole wash-out to anaphase onset) in RCC1-positive control (68.7 ± 2.1, n=46) and RCC1-depleted cells (61.5 ± 1.4, n=47). Bars indicate means ± SDs from >3 independent experiments. * indicates statistical significance according to Welch’s *t*-test (p = 0.0061). Scale bars = 10 μm.

**Figure S3.**
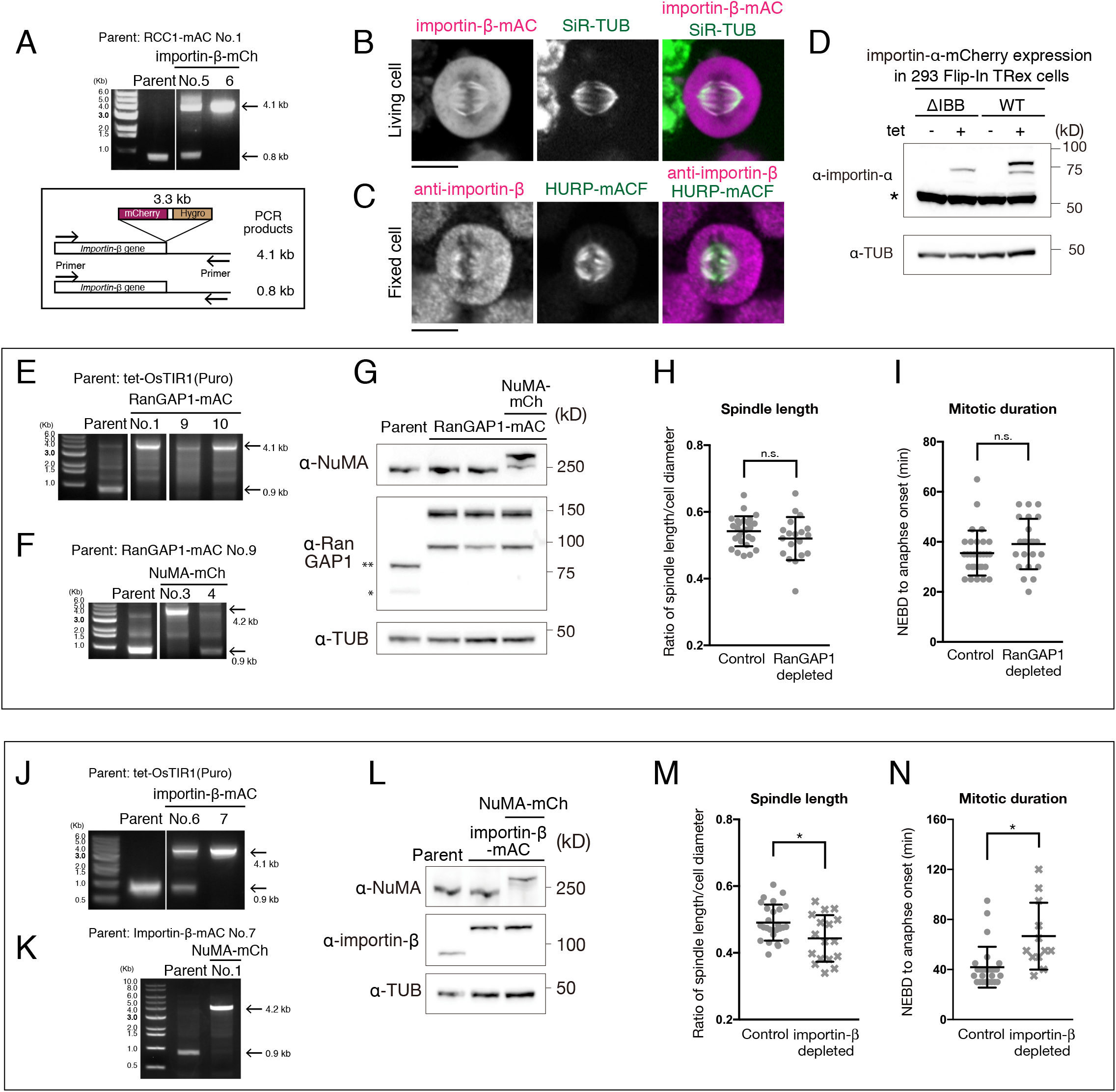
Generation of cell lines for auxin-inducible degradation of endogenous Ran-GAP1 and importin-β. (A) Genomic PCR showing clone genotypes after hygromycin (Hygro) selection. Clones No.6 were used. (B) Metaphase importin-β-mAC cells showing live fluorescent images of importin-β-mAC, and SiR-TUB. Single z-section images are shown. (C) Immunofluorescence images of fixed metaphase cells showing k-fiber localization endogenous importin-β and mAID-tagged HURP (HURP-mACF). Maximally projected images from 3 z-sections are shown. (D) Immunoblotting for anti-importin-α and anti-α-tubulin (TUB, loading control) showing ectopic expression of the importin-α wild type (WT, right) and a ΔIBB mutant (left) following Dox treatment. * indicates endogenous importin-α. (E) Genomic PCR showing clone genotypes after neomycin (Neo) selection. Clone No.9 was used as a parental cell in the second selections. (F) Genomic PCR showing clone genotypes after hygromycin (Hygro) selection. Clone No.3 (NuMA-mCh) was selected. (G) Immunoblotting for anti-NuMA, anti-RanGAP1 and anti-α-tubulin (TUB, loading control) showing bi-allelic insertion of the indicated tags. * and ** indicate RanGAP1 and SUMO-1 conjugated RanGAP1, respectively. (H) Scatterplots of the ratio of spindle length and cell diameter in control (0.54 ± 0.04, n=26) and RanGAP1-depleted (0.52 ± 0.07, n=19) cells. (I) Scatterplots of mitotic duration (NEBD to anaphase onset) in control (35.5 ± 9.0, n=29) and RanGAP1-depleted (39.1 ± 10.1, n=23) cells. Bars in (H) and (I) indicate means ± SDs from >3 independent experiments. Differences were not statistically significant based on Welch’s *t*-test in H (p=0.2108) and I (p=0.1851). (J) Genomic PCR showing clone genotypes after neomycin (Neo) selection. Clone No.7 was used as a parental cell in the second selections. (K) Genomic PCR showing clone genotype after hygromycin (Hygro) selection. Clone No.1 was selected for further use. (L) Western blot detection using anti-NuMA, anti-importin-β and anti-α-tubulin antibodies (TUB, loading control) showing bi-allelic insertion of the indicated tags. (M) Scatterplots of the ratio of spindle length and cell diameter in control (0.49 ± 0,05, n=26) and importin-β-depleted (0.44 ± 0.07, n=17) cells. (N) Scatterplots of mitotic duration (NEBD to anaphase onset) in control (41.9 ± 16.3, n=27) and importin-β-depleted (66.7 ± 26.7, n=12) cells. Bars in (M) and (N) indicate mean ± SD from >3 independent experiments. * indicates statistical significance according to Welch’s *t*-test (p<0.05) in (M) and (N). Scale bars = 10 μm.

**Figure S4.**
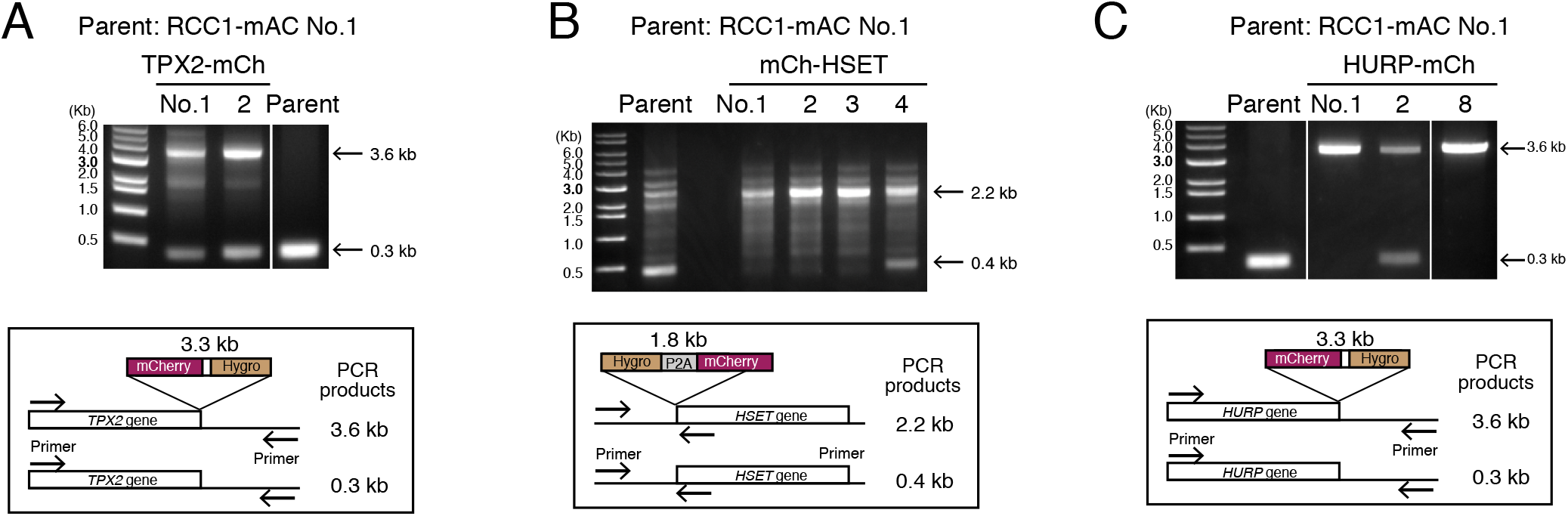
Generation of double knock-in cell lines that express RCC1-mAC and mCherry-fused TPX2, HSET, or HURP. (A-C) Genomic PCR showing clone genotypes after hygromycin (Hygro) selections. Clones No.1 (A), No. 1 (B), and No.8 (C) were used. The mCherry cassette was inserted into only one copy of TPX2 gene loci (A).

**Figure S5.**
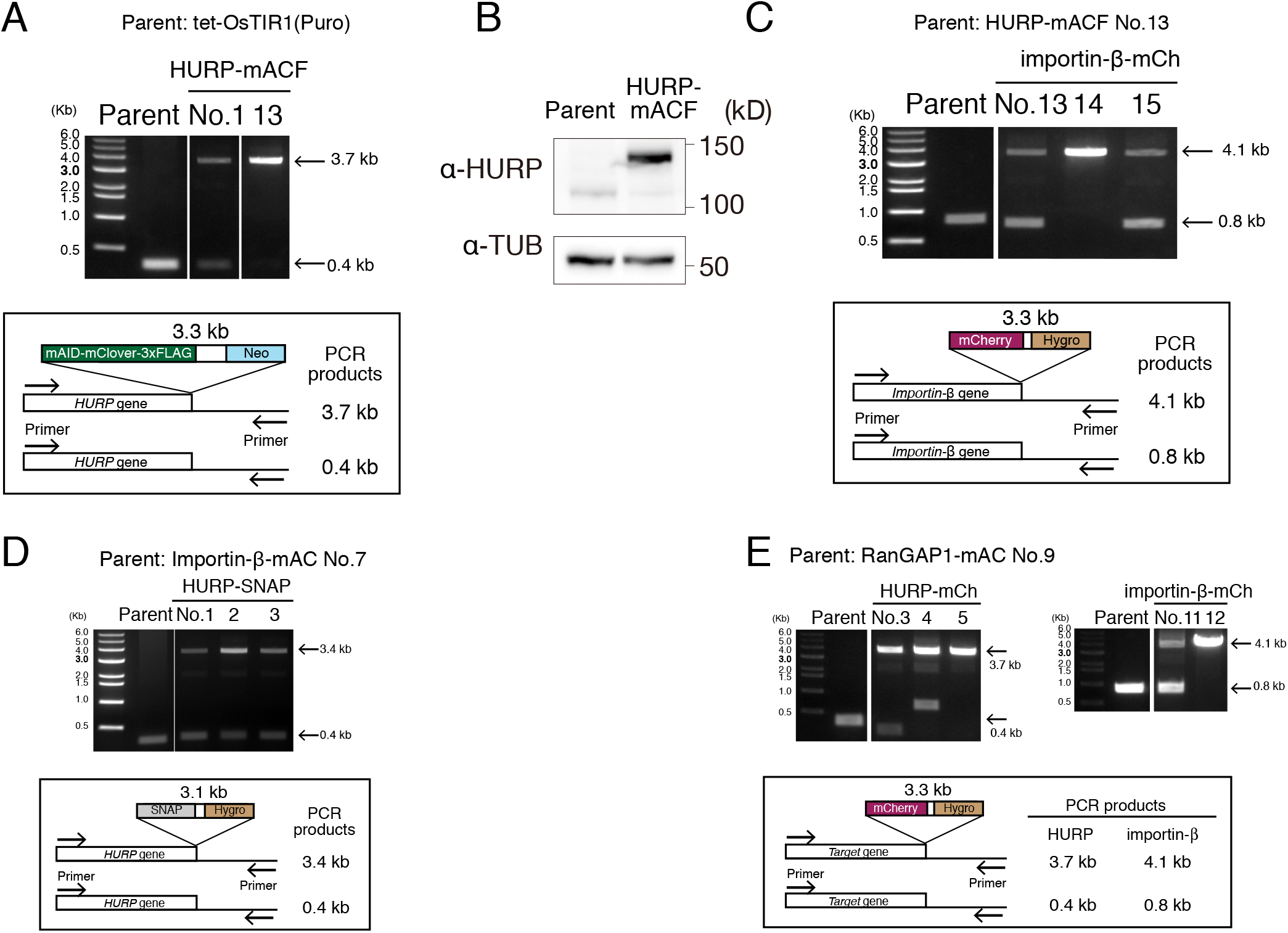
Generation of cell lines that degrade or visualize endogenous HURP. (A) Genomic PCR showing the clone genotype after neomycin (Neo) selection. Clone No.13 was used as a parental cell in subsequent selections. (B) Immunoblotting for anti-HURP and anti-α-tubulin (TUB, loading control) showing bi-allelic insertion of the indicated tags. (C-E) Genomic PCR showing clone genotypes after hygromycin (Hygro) selection. Clone No.14 (C), No. 3 (D), No. 5 (E: HURP-mCh), and No. 12 (E: importin-β) were used, respectively. The SNAP cassette was inserted into only one copy of HURP gene loci (D).

**Figure S6.**
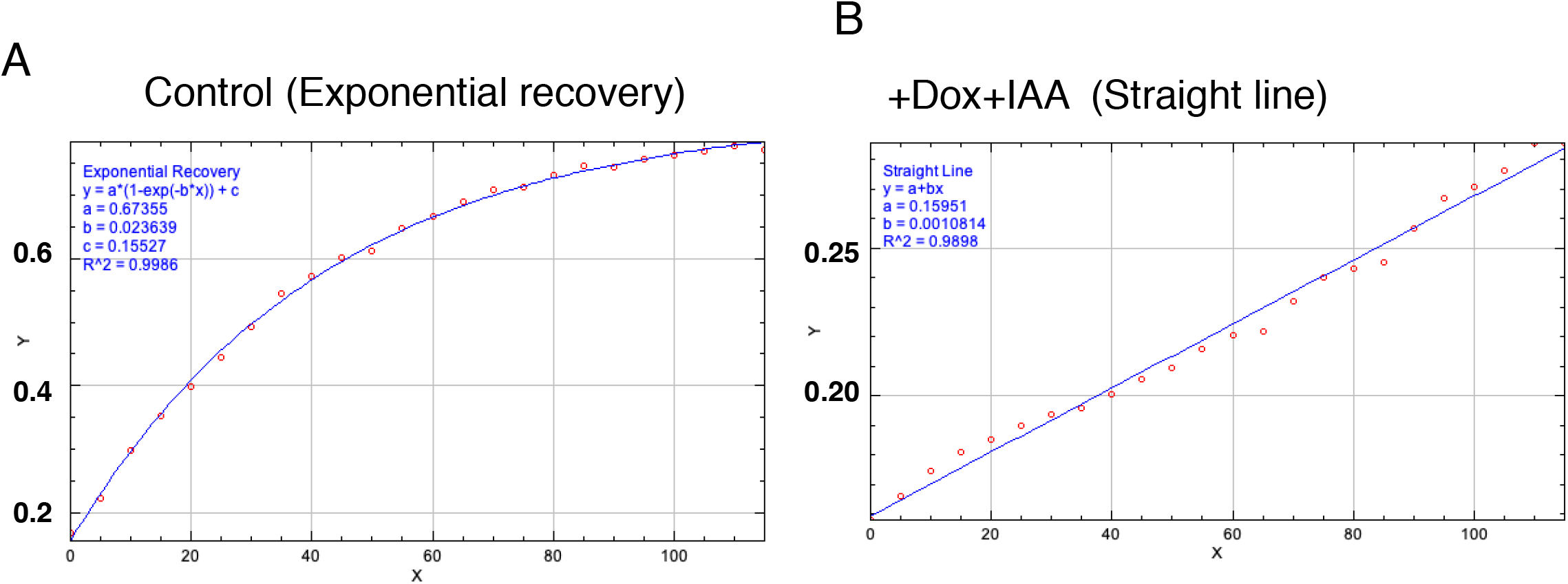
Fluorescent recovery kinetics of HURP in the presence or absence of importin-β. (A-B) Graphs showing a fitted curve or a straight line on each plot. Formulas and parameters are also indicated.

**Table S1:** Cell lines used in this study.

| No. | Name | Description | Clo<br>ne<br>No. | Plasmids used | Par<br>ent<br>al<br>cell | Reference |
| --- | --- | --- | --- | --- | --- | --- |
| 1 | HCT116 tet-OsTIR1 | AAVS1::PTRE3G OsTIR1 (Puro) |  | pAAVS1 T2 and<br>MK243<br>(Addgene#7283<br>5) |  | [35] |
| 2 | NuMA-mACF +<br>DHC-SNAP +<br>mCh-NuMA WT | AAVS1::PTRE3G OsTIR1 (Puro), NuMA1::<br>NuMA-mAID-mClover-3FLAG (Neo), DHC1::<br>DHC-SNAP (BSD), Rosa26:: PTRE3G<br>mCherry-NuMA WT (Hygro) | 7 | hROSA26<br>CRISPR-pX330<br>and pTK503 |  | [20] |
| 3 | NuMA-mACF +<br>DHC-SNAP +<br>mCh-NuMA ΔNLS | AAVS1::PTRE3G OsTIR1 (Puro), NuMA1::<br>NuMA-mAID-mClover-3FLAG (Neo), DHC1::<br>DHC-SNAP (BSD), Rosa26:: PTRE3G<br>mCherry-NuMA ΔNLS (Hygro) | 1 | hROSA26<br>CRISPR-pX330<br>and pTK699 | 2 | This study |
| 4 | NuMA-mACF +<br>DHC-SNAP +<br>mCh-NuMAΔex24 | AAVS1::PTRE3G OsTIR1 (Puro), NuMA1::<br>NuMA-mAID-mClover-3FLAG (Neo), DHC1::<br>DHC-SNAP (BSD), Rosa26:: PTRE3G<br>mCherry-NuMA Δex24 (Hygro) | 1 | hROSA26<br>CRISPR-pX330<br>and pTK700 | 2 | This study |
| 5 | NuMA-mACF +<br>DHC-SNAP +<br>mCh-NuMA<br>$\Delta$(NLS+MTBD2) | AAVS1::PTRE3G OsTIR1 (Puro), NuMA1::<br>NuMA-mAID-mClover-3FLAG (Neo), DHC1::<br>DHC-SNAP (BSD), Rosa26:: PTRE3G<br>mCherry-NuMA $\Delta$(NLS+MTBD2) (Hygro) | 1 | hROSA26<br>CRISPR-pX330<br>and pTK509 | 2 | This study |
| 6 | NuMA-mACF +<br>DHC-SNAP +<br>mCh-NuMA ΔC-ter | AAVS1::PTRE3G OsTIR1 (Puro), NuMA1::<br>NuMA-mAID-mClover-3FLAG (Neo), DHC1::<br>DHC-SNAP (BSD), Rosa26:: PTRE3G<br>mCherry-NuMA ΔC-ter (Hygro) | 3 | hROSA26<br>CRISPR-pX330<br>and pTK510 | 2 | This study |
| 7 | Flip-In T-REx 293 | Invitrogen |  |  |  | [57] |
| 8 | Flip-In T-REx 293<br>importin-α-WT<br>mCherry | Flip-In:: importin-α-WT mCherry (hygro) | Poly<br>clon<br>al | pOG44 and<br>pTK960 | 7 | This study |
| 9 | Flip-In T-REx 293<br>importin-α-ΔIBB<br>mCherry | Flip-In:: importin-α-ΔIBB mCherry (hygro) | Poly<br>clon<br>al | pOG44 and<br>pTK961 | 7 | This study |
| 10 | RCC1-mAC | AAVS1::PTRE3G OsTIR1 (Puro), RCC1::<br>RCC1-mAID-mClover (Neo) | 1 | pTK361+<br>pHH45 | 1 | This study |
| 11 | RCC1-mAC +<br>NuMA-mCh | AAVS1::PTRE3G OsTIR1 (Puro), RCC1::<br>RCC1-mAID-mClover (Neo), NuMA1::<br>NuMA-mCh (Hygro) | 1 | pTK372+<br>pTK435 | 10 | This study |
| 12 | RanGAP1-mAC | AAVS1::PTRE3G OsTIR1 (Puro),<br>RanGAP1:: RanGAP1-mAID-mClover (Neo) | 9 | pHH49 +<br>pHH51 | 1 | This study |
| 13 | RanGAP1-mAC +<br>NuMA-mCh | AAVS1::PTRE3G OsTIR1 (Puro),<br>RanGAP1:: RanGAP1-mAID-mClover (Neo),<br>NuMA1:: NuMA-mCh (Hygro) | 5 | pTK372+<br>pTK435 | 12 | This study |
| 14 | importin-β-mAC | AAVS1::PTRE3G OsTIR1 (Puro), importin-<br>$\beta$:: importin-β-mAID-mClover (Neo) | 7 | pHH50 +<br>pHH57 | 1 | This study |
| 15 | importin-β-mAC +<br>NuMA-mCh | AAVS1::PTRE3G OsTIR1 (Puro), importin-<br>$\beta$:: importin-β-mAID-mClover (Neo),<br>NuMA1:: NuMA-mCh (Hygro) | 1 | pTK372+<br>pTK435 | 14 | This study |
| 16 | RCC1-mAC +<br>importin-β-mCh | AAVS1::PTRE3G OsTIR1 (Puro), RCC1::<br>RCC1-mAID-mClover (Neo), NuMA1::<br>NuMA-mCh (Hygro) | 6 | pHH50 +<br>pTK481 | 10 | This study |
| 17 | RCC1-mAC + HURP-mCh | AAVS1::PTRE3G OsTIR1 (Puro), RCC1::RCC1-mAID-mClover (Neo), HURP::HURP-mCh (Hygro) | 8 | pTK532+ pTK541 | 10 | This study |
| 18 | RCC1-mAC + TPX2-mCh | AAVS1::PTRE3G OsTIR1 (Puro), RCC1::RCC1-mAID-mClover (Neo), TPX2::TPX2-mCh (Hygro) | 1 | pTK527+ pTK502 | 10 | This study |
| 19 | RCC1-mAC + mCh-HSET | AAVS1::PTRE3G OsTIR1 (Puro), RCC1::RCC1-mAID-mClover (Neo), HSET::mCh-HSET (Hygro) | 1 | pTK523+ pTK531 | 10 | This study |
| 20 | RanGAP1-mAC + HURP-mCh | AAVS1::PTRE3G OsTIR1 (Puro), RanGAP1::RanGAP1-mAID-mClover (Neo), HURP::HURP-mCh (Hygro) | 5 | pTK532+ pTK541 | 12 | This study |
| 21 | RanGAP1-mAC + importin-β-mCh | AAVS1::PTRE3G OsTIR1 (Puro), RanGAP1::RanGAP1-mAID-mClover (Neo), importin-β::importin-β-mCh (Hygro) | 12 | pHH50 + pTK481 | 12 | This study |
| 22 | importin-β-mAC + HURP-SNAP | AAVS1::PTRE3G OsTIR1 (Puro), importin-β::importin-β-mAID-mClover (Neo), HURP::HURP-SNAP (Hygro) | 3 | pTK532+ pTK589 | 14 | This study |
| 23 | HURP-mACF | AAVS1::PTRE3G OsTIR1 (Puro), HURP::HURP-mAID-mClover-3FLAG (Neo) | 13 | pTK532+ pTK596 | 1 | This study |
| 24 | HURP-mACF + importin-β-mCh | AAVS1::PTRE3G OsTIR1 (Puro), HURP::HURP-mAID-mClover-3FLAG (Neo), importin-β::importin-β-mCh (Hygro) | 14 | pHH50 + pTK481 | 23 | This study |

**Table S2:**
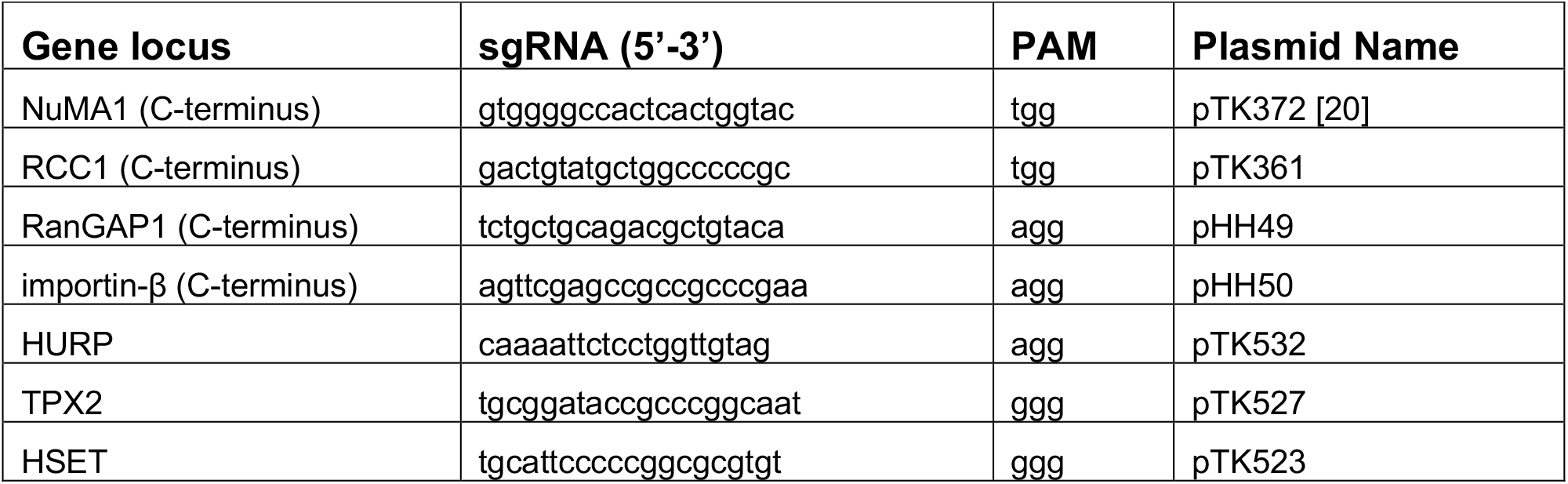
sgRNA sequences for CRISPR/Cas9-mediated genome editing

**Table S3:**
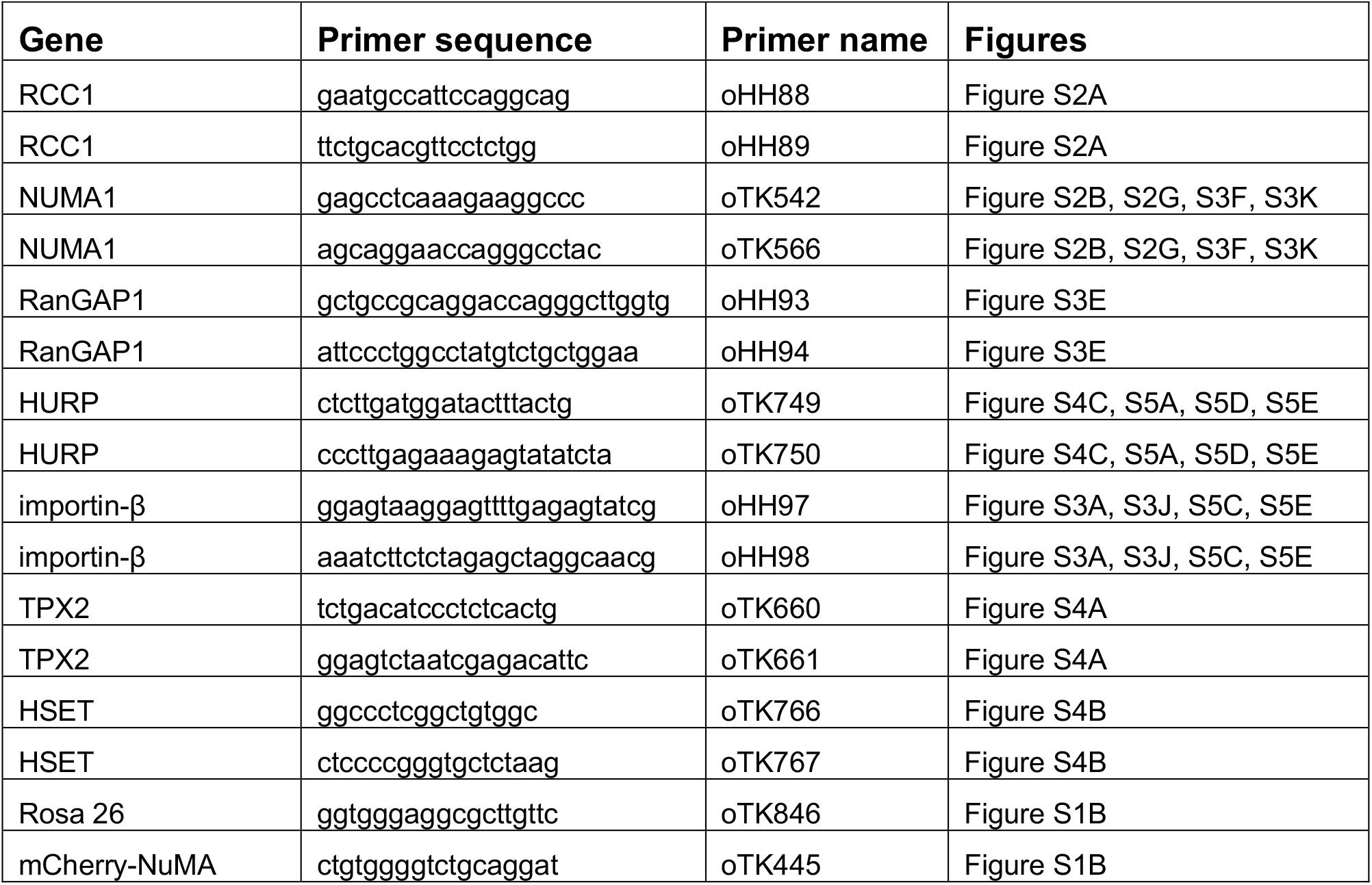
PCR primers used to confirm gene editing

## Notes

### Competing Interest Statement

The authors have declared no competing interest.

## References

1. Reber, S., and Hyman, A.A. (2015). Emergent Properties of the Metaphase Spindle. Cold Spring Harb Perspect Biol 7, a015784.

2. Heald, R., and Khodjakov, A. (2015). Thirty years of search and capture: The complex simplicity of mitotic spindle assembly. J Cell Biol 211, 1103–1111.

3. Kalab, P., and Heald, R. (2008). The RanGTP gradient - a GPS for the mitotic spindle. J Cell Sci 121, 1577–1586.

4. Forbes, D.J., Travesa, A., Nord, M.S., and Bernis, C. (2015). Reprint of “Nuclear transport factors: global regulation of mitosis”. Curr Opin Cell Biol 34, 122–134.

5. Bischoff, F.R., and Ponstingl, H. (1991). Catalysis of guanine nucleotide exchange on Ran by the mitotic regulator RCC1. Nature 354, 80–82.

6. Bischoff, F.R., Klebe, C., Kretschmer, J., Wittinghofer, A., and Ponstingl, H. (1994). RanGAP1 induces GTPase activity of nuclear Ras-related Ran. Proc Natl Acad Sci U S A 91, 2587–2591.

7. Kalab, P., Weis, K., and Heald, R. (2002). Visualization of a Ran-GTP gradient in interphase and mitotic Xenopus egg extracts. Science 295, 2452–2456.

8. Kalab, P., Pralle, A., Isacoff, E.Y., Heald, R., and Weis, K. (2006). Analysis of a RanGTP-regulated gradient in mitotic somatic cells. Nature 440, 697–701.

9. Dumont, J., Petri, S., Pellegrin, F., Terret, M.E., Bohnsack, M.T., Rassinier, P., Georget, V., Kalab, P., Gruss, O.J., and Verlhac, M.H. (2007). A centriole- and RanGTP-independent spindle assembly pathway in meiosis I of vertebrate oocytes. J Cell Biol 176, 295–305.

10. Moutinho-Pereira, S., Stuurman, N., Afonso, O., Hornsveld, M., Aguiar, P., Goshima, G., Vale, R.D., and Maiato, H. (2013). Genes involved in centrosome-independent mitotic spindle assembly in Drosophila S2 cells. Proc Natl Acad Sci U S A 110, 19808–19813.

11. Hasegawa, K., Ryu, S.J., and Kalab, P. (2013). Chromosomal gain promotes formation of a steep RanGTP gradient that drives mitosis in aneuploid cells. J Cell Biol 200, 151–161.

12. Holubcova, Z., Blayney, M., Elder, K., and Schuh, M. (2015). Human oocytes. Error-prone chromosome-mediated spindle assembly favors chromosome segregation defects in human oocytes. Science 348, 1143–1147.

13. Drutovic, D., Duan, X., Li, R., Kalab, P., and Solc, P. (2020). RanGTP and importin beta regulate meiosis I spindle assembly and function in mouse oocytes. EMBO J 39, e101689.

14. Furuta, M., Hori, T., and Fukagawa, T. (2016). Chromatin binding of RCC1 during mitosis is important for its nuclear localization in interphase. Mol Biol Cell 27, 371–381.

15. Nachury, M.V., Maresca, T.J., Salmon, W.C., Waterman-Storer, C.M., Heald, R., and Weis, K. (2001). Importin beta is a mitotic target of the small GTPase Ran in spindle assembly. Cell 104, 95–106.

16. Wiese, C., Wilde, A., Moore, M.S., Adam, S.A., Merdes, A., and Zheng, Y. (2001). Role of importin-beta in coupling Ran to downstream targets in microtubule assembly. Science 291, 653–656.

17. Gruss, O.J., Carazo-Salas, R.E., Schatz, C.A., Guarguaglini, G., Kast, J., Wilm, M., Le Bot, N., Vernos, I., Karsenti, E., and Mattaj, I.W. (2001). Ran induces spindle assembly by reversing the inhibitory effect of importin alpha on TPX2 activity. Cell 104, 83–93.

18. Stewart, M. (2007). Molecular mechanism of the nuclear protein import cycle. Nat Rev Mol Cell Biol 8, 195–208.

19. Hueschen, C.L., Kenny, S.J., Xu, K., and Dumont, S. (2017). NuMA recruits dynein activity to microtubule minus-ends at mitosis. Elife 6.

20. Okumura, M., Natsume, T., Kanemaki, M.T., and Kiyomitsu, T. (2018). Dynein-Dynactin-NuMA clusters generate cortical spindle-pulling forces as a multi-arm ensemble. Elife 7.

21. Gaglio, T., Saredi, A., and Compton, D.A. (1995). NuMA is required for the organization of microtubules into aster-like mitotic arrays. J Cell Biol 131, 693–708.

22. Silk, A.D., Holland, A.J., and Cleveland, D.W. (2009). Requirements for NuMA in maintenance and establishment of mammalian spindle poles. J Cell Biol 184, 677–690.

23. Wittmann, T., Wilm, M., Karsenti, E., and Vernos, I. (2000). TPX2, A novel xenopus MAP involved in spindle pole organization. J Cell Biol 149, 1405–1418.

24. Garrett, S., Auer, K., Compton, D.A., and Kapoor, T.M. (2002). hTPX2 is required for normal spindle morphology and centrosome integrity during vertebrate cell division. Curr Biol 12, 2055–2059.

25. Petry, S., Groen, A.C., Ishihara, K., Mitchison, T.J., and Vale, R.D. (2013). Branching microtubule nucleation in Xenopus egg extracts mediated by augmin and TPX2. Cell 152, 768–777.

26. Roostalu, J., Cade, N.I., and Surrey, T. (2015). Complementary activities of TPX2 and chTOG constitute an efficient importin-regulated microtubule nucleation module. Nat Cell Biol 17, 1422–1434.

27. King, M.R., and Petry, S. (2020). Phase separation of TPX2 enhances and spatially coordinates microtubule nucleation. Nat Commun 11, 270.

28. Ems-McClung, S.C., Zheng, Y., and Walczak, C.E. (2004). Importin alpha/beta and Ran-GTP regulate XCTK2 microtubule binding through a bipartite nuclear localization signal. Mol Biol Cell 15, 46–57.

29. Ems-McClung, S.C., Emch, M., Zhang, S., Mahnoor, S., Weaver, L.N., and Walczak, C.E. (2020). RanGTP induces an effector gradient of XCTK2 and importin alpha/beta for spindle microtubule cross-linking. J Cell Biol 219.

30. Cai, S., Weaver, L.N., Ems-McClung, S.C., and Walczak, C.E. (2009). Kinesin-14 family proteins HSET/XCTK2 control spindle length by cross-linking and sliding microtubules. Mol Biol Cell 20, 1348-1359.

31. Sillje, H.H., Nagel, S., Korner, R., and Nigg, E.A. (2006). HURP is a Ran-importin beta-regulated protein that stabilizes kinetochore microtubules in the vicinity of chromosomes. Curr Biol 16, 731–742.

32. Chang, C.C., Huang, T.L., Shimamoto, Y., Tsai, S.Y., and Hsia, K.C. (2017). Regulation of mitotic spindle assembly factor NuMA by Importin-beta. J Cell Biol 216, 3453–3462.

33. Giesecke, A., and Stewart, M. (2010). Novel binding of the mitotic regulator TPX2 (target protein for Xenopus kinesin-like protein 2) to importin-alpha. J Biol Chem 285, 17628–17635.

34. Kiyomitsu, T. (2019). The cortical force-generating machinery: how cortical spindle-pulling forces are generated. Curr Opin Cell Biol 60, 1–8.

35. Natsume, T., Kiyomitsu, T., Saga, Y., and Kanemaki, M.T. (2016). Rapid Protein Depletion in Human Cells by Auxin-Inducible Degron Tagging with Short Homology Donors. Cell Rep 15, 210–218.

36. Seldin, L., Muroyama, A., and Lechler, T. (2016). NuMA-microtubule interactions are critical for spindle orientation and the morphogenesis of diverse epidermal structures. Elife 5.

37. Gallini, S., Carminati, M., De Mattia, F., Pirovano, L., Martini, E., Oldani, A., Asteriti, I.A., Guarguaglini, G., and Mapelli, M. (2016). NuMA Phosphorylation by Aurora-A Orchestrates Spindle Orientation. Curr Biol 26, 458–469.

38. Siller, K.H., Cabernard, C., and Doe, C.Q. (2006). The NuMA-related Mud protein binds Pins and regulates spindle orientation in Drosophila neuroblasts. Nat Cell Biol 8, 594–600.

39. Du, Q., Stukenberg, P.T., and Macara, I.G. (2001). A mammalian Partner of inscuteable binds NuMA and regulates mitotic spindle organization. Nat Cell Biol 3, 1069–1075.

40. Tang, T.K., Tang, C.J., Chao, Y.J., and Wu, C.W. (1994). Nuclear mitotic apparatus protein (NuMA): spindle association, nuclear targeting and differential subcellular localization of various NuMA isoforms. J Cell Sci 107 (Pt 6), 1389–1402.

41. Kohler, M., Speck, C., Christiansen, M., Bischoff, F.R., Prehn, S., Haller, H., Gorlich, D., and Hartmann, E. (1999). Evidence for distinct substrate specificities of importin alpha family members in nuclear protein import. Mol Cell Biol 19, 7782–7791.

42. Ciciarello, M., Mangiacasale, R., Thibier, C., Guarguaglini, G., Marchetti, E., Di Fiore, B., and Lavia, P. (2004). Importin beta is transported to spindle poles during mitosis and regulates Ran-dependent spindle assembly factors in mammalian cells. J Cell Sci 117, 6511–6522.

43. Song, L., Craney, A., and Rape, M. (2014). Microtubule-dependent regulation of mitotic protein degradation. Mol Cell 53, 179–192.

44. Sackton, K.L., Dimova, N., Zeng, X., Tian, W., Zhang, M., Sackton, T.B., Meaders, J., Pfaff, K.L., Sigoillot, F., Yu, H., et al. (2014). Synergistic blockade of mitotic exit by two chemical inhibitors of the APC/C. Nature 514, 646–649.

45. Wei, J.H., Zhang, Z.C., Wynn, R.M., and Seemann, J. (2015). GM130 Regulates Golgi-Derived Spindle Assembly by Activating TPX2 and Capturing Microtubules. Cell 162, 287–299.

46. Brownlee, C., and Heald, R. (2019). Importin alpha Partitioning to the Plasma Membrane Regulates Intracellular Scaling. Cell 176, 805–815 e808.

47. Chinen, T., Yamamoto, S., Takeda, Y., Watanabe, K., Kuroki, K., Hashimoto, K., Takao, D., and Kitagawa, D. (2020). NuMA assemblies organize microtubule asters to establish spindle bipolarity in acentrosomal human cells. EMBO J 39, e102378.

48. Lorson, M.A., Horvitz, H.R., and van den Heuvel, S. (2000). LIN-5 is a novel component of the spindle apparatus required for chromosome segregation and cleavage plane specification in Caenorhabditis elegans. J Cell Biol 148, 73–86.

49. Greenberg, S.R., Tan, W., and Lee, W.L. (2018). Num1 versus NuMA: insights from two functionally homologous proteins. Biophys Rev 10, 1631–1636.

50. Weaver, L.N., Ems-McClung, S.C., Chen, S.H., Yang, G., Shaw, S.L., and Walczak, C.E. (2015). The Ran-GTP gradient spatially regulates XCTK2 in the spindle. Curr Biol 25, 1509–1514.

51. Sikirzhytski, V., Renda, F., Tikhonenko, I., Magidson, V., McEwen, B.F., and Khodjakov, A. (2018). Microtubules assemble near most kinetochores during early prometaphase in human cells. J Cell Biol 217, 2647–2659.

52. Booth, D.G., Hood, F.E., Prior, I.A., and Royle, S.J. (2011). A TACC3/ch-TOG/clathrin complex stabilises kinetochore fibres by inter-microtubule bridging. EMBO J 30, 906–919.

53. Nishimoto, T., Eilen, E., and Basilico, C. (1978). Premature of chromosome condensation in a ts DNA-mutant of BHK cells. Cell 15, 475–483.

54. Kiyomitsu, T., and Cheeseman, I.M. (2012). Chromosome-and spindle-pole-derived signals generate an intrinsic code for spindle position and orientation. Nat Cell Biol 14, 311–317.

55. Soderholm, J.F., Bird, S.L., Kalab, P., Sampathkumar, Y., Hasegawa, K., Uehara-Bingen, M., Weis, K., and Heald, R. (2011). Importazole, a small molecule inhibitor of the transport receptor importin-beta. ACS Chem Biol 6, 700–708.

56. Kiyomitsu, T., and Cheeseman, I.M. (2013). Cortical dynein and asymmetric membrane elongation coordinately position the spindle in anaphase. Cell 154, 391–402.

57. Kiyomitsu, T., Murakami, H., and Yanagida, M. (2011). Protein interaction domain mapping of human kinetochore protein Blinkin reveals a consensus motif for binding of spindle assembly checkpoint proteins Bub1 and BubR1. Mol Cell Biol 31, 998–1011.

58. Goshima, G., Nedelec, F., and Vale, R.D. (2005). Mechanisms for focusing mitotic spindle poles by minus end-directed motor proteins. J Cell Biol 171, 229–240.

